# Maximizing Heterologous Expression of Engineered Type I Polyketide Synthases: Investigating Codon Optimization Strategies

**DOI:** 10.1101/2023.06.13.544731

**Authors:** Matthias Schmidt, Namil Lee, Chunjun Zhan, Jacob B. Roberts, Alberto A. Nava, Leah Keiser, Aaron Vilchez, Yan Chen, Christopher J. Petzold, Robert W. Haushalter, Lars M. Blank, Jay D. Keasling

## Abstract

Type I polyketide synthases (T1PKSs) hold an enormous potential as a rational production platform for the biosynthesis of specialty chemicals. However, despite the great progress in this field, the heterologous expression of PKSs remains a major challenge. One of the first measures to improve heterologous gene expression can be codon optimization. Although controversial, choosing the wrong codon optimization strategy can have detrimental effects on protein and product levels. In this study, we analyzed 11 different codon variants of an engineered T1PKS and investigated in a systematic approach their influence on heterologous expression in *Corynebacterium glutamicum*, *Escherichia coli*, and *Pseudomonas putida*. Our best performing codon variants exhibited a minimum 50-fold increase in PKS protein levels, which also enables the production of an unnatural polyketide in each of the hosts. Furthermore, we developed a free online tool (https://basebuddy.lbl.gov) that offers transparent and highly customizable codon optimization with up-to-date codon usage tables.

Here, we not only highlight the significance of codon optimization but also establish the groundwork for high-throughput assembly and characterization of PKS pathways in alternative hosts.

## 1 INTRODUCTION

Type I polyketide synthases (T1PKSs) are a class of natural enzymes primarily found in bacteria or fungi that are responsible for the biosynthesis of secondary metabolites. Over the years, humans have harnessed the therapeutic potential of these metabolites, and numerous indispensable drugs have been discovered (1).

The modular architecture of T1PKSs enables the iterative assembly of long carbon chains, while also providing the flexibility for the optional incorporation of functional groups (2). The theoretical design space offered by this modularity has attracted significant research attention, and T1PKSs have been successfully reprogrammed for the production of unnatural polyketides (3–5).

However, one of the biggest challenges in PKS engineering is the native host itself. Most of the discovered PKSs originate from the genus *Streptomyces*, a GC-rich, gram-positive and filamentous bacterium. While certain streptomycetes have been highly domesticated and optimized for the production of larger, high value molecules, their efficacy for the production of industrially relevant bulk chemicals is limited (6). To date, little progress has been made in exploring alternative PKS hosts. Besides the required genetic modifications for PKS expression, polyketide titers in non-native hosts are often very low, which can usually be traced back to poor precursor availability or low protein levels (7, 8). Common strategies for improving polyketide titers include the supplementation of the media, the exchange of promoters, or codon optimization (4). However, the importance of the latter one is often neglected or underestimated.

The host’s codon preference usually has a distinct pattern and can be summarized in codon usage tables (9). The choice of specific codons can impact transcription and translation rates, and is also involved in expression control mechanisms. Well-studied examples include the rare *Streptomyces* codon TTA and the use of alternative start codons such as GTG or TTG (10, 11).

Codon optimization represents a strategy to address these variations in codon preferences during heterologous gene expression. This approach involves selectively substituting specific codons while preserving the amino acid sequence of the protein. There are three commonly used strategies for this purpose: (i) replacing the original codons with the most frequently used codon of the targeted host (12), (ii) matching the codon frequency of the targeted host (13) and (iii) harmonizing the codon frequency of the targeted host with the codon frequency of the native host (14). After codon optimization, the final nucleotide sequence will likely be significantly different from the original sequence, and many researchers have to rely on DNA synthesis services to synthesize the codon-optimized gene. Although convenient, the optimization algorithms offered by most synthesis services are not publicly available and their functionality is very limited. The recently published open-source codon optimization tool DNA Chisel offers an easy way to apply these aforementioned methods and further customize the resulting nucleotide sequence (15).

In this study, we investigated the expression and activity of an engineered T1PKS with different codon variants in the three hosts *Corynebacterium glutamicum, Escherichia coli*, and *Pseudomonas putida*. *E. coli* and *C. glutamicum* are well-established industrial hosts for the large-scale production of proteins and small molecules, while *P. putida* has shown enormous potential for the valorization of renewable feedstocks (16–18). By targeting three heterologous hosts and applying the three most common gene optimizations methods, we designed 9 codon variants of the engineered PKS (Figure 1). Furthermore, as a conventional approach, we obtained and cloned the native sequences from *Streptomyces aureofaciens* Tü117 and *Saccharopolyspora erythraea* NRRL2338, and also tested the effect of two different start codons, GTG and ATG. Due to the large size of PKSs and the metabolic burden of plasmid maintenance, we developed a backbone excision-dependent expression (BEDEX) system to facilitate the cloning process and enable constitutive expression in the heterologous hosts. Also, we hypothesized that the strong repression of BEDEX vectors would decrease the occurrence of mutations during DNA assembly. We further confirmed the universal functionality of BEDEX vectors in our selected hosts and applied the BEDEX system to heterologously express the 11 codon variants of the engineered PKS. To characterize our codon variants *in vivo*, we measured the PKS protein and transcript levels and demonstrated the production of an unnatural polyketide in each host.

**Figure 1:**
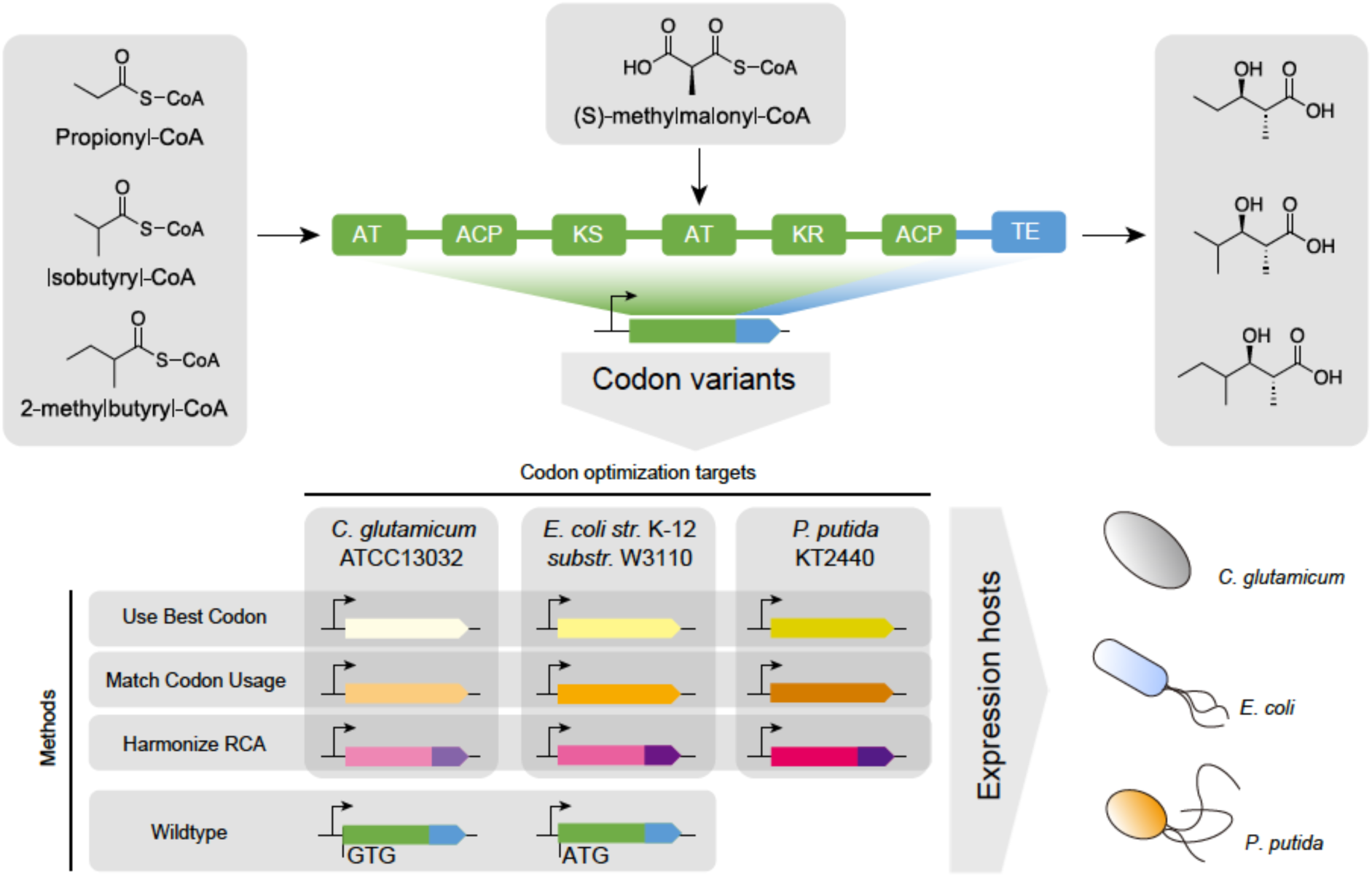
Engineered polyketide synthase with applied codon optimization strategies and targeted heterologous hosts. The loading module and module 1 originates from the lipomycin polyketide synthase (LipPKS) from *Streptomyces aureofaciens* Tü117 (green). The thioesterase domain (TE) originates from the erythromycin PKS (EryPKS) from *Saccharopolyspora erythraea* NRRL2338 (blue). By fusing these two parts together, the engineered PKS design yields a variety of short-chain 3-hydroxy acids. The gene sequence of the reprogrammed LipPKS was codon optimized using the DNA Chisel algorithms for ‘Use Best Codon’ (yellow), ‘Match Codon Usage’ (orange) and ‘Harmonize RCA’ (red/purple). The ‘Harmonize RCA’ algorithm required to codon optimize the LipPKS and EryPKS parts separately. Codon optimizations targeted the three heterologous hosts *C. glutamicum, E. coli*, and *P. putida*.

Optimizing the gene sequence for heterologous expression is a controversial topic and systematic studies are scarce, especially in the field of PKS engineering. Here, we showcase the importance of codon optimization and lay the foundation for high-throughput assembly and PKS pathway discovery.

## 2 RESULTS

### 2.1 Comparative analysis on codon usage patterns of host organisms

To conduct a comprehensive analysis of the overall codon usage in the target host organisms, we compared the codon usage patterns among three native PKS hosts, our selected heterologous host species, and other well-studied organisms as outgroup references. 16S rRNA sequence-based phylogenetic analysis on these species shows that *C. glutamicum* is the most closely related species to *Streptomyces*, and that *E. coli* and *P. putida* are more closely related to each other (Figure 2a). Based on the phylogenetic data, it could be concluded that *C. glutamicum* should be the most suitable heterologous host for genes that are sourced from streptomycetes. However, their codon usage might be vastly different since the GC-content in *Streptomyces* is about 20 % higher than in *C. glutamicum*.

**Figure 2:**
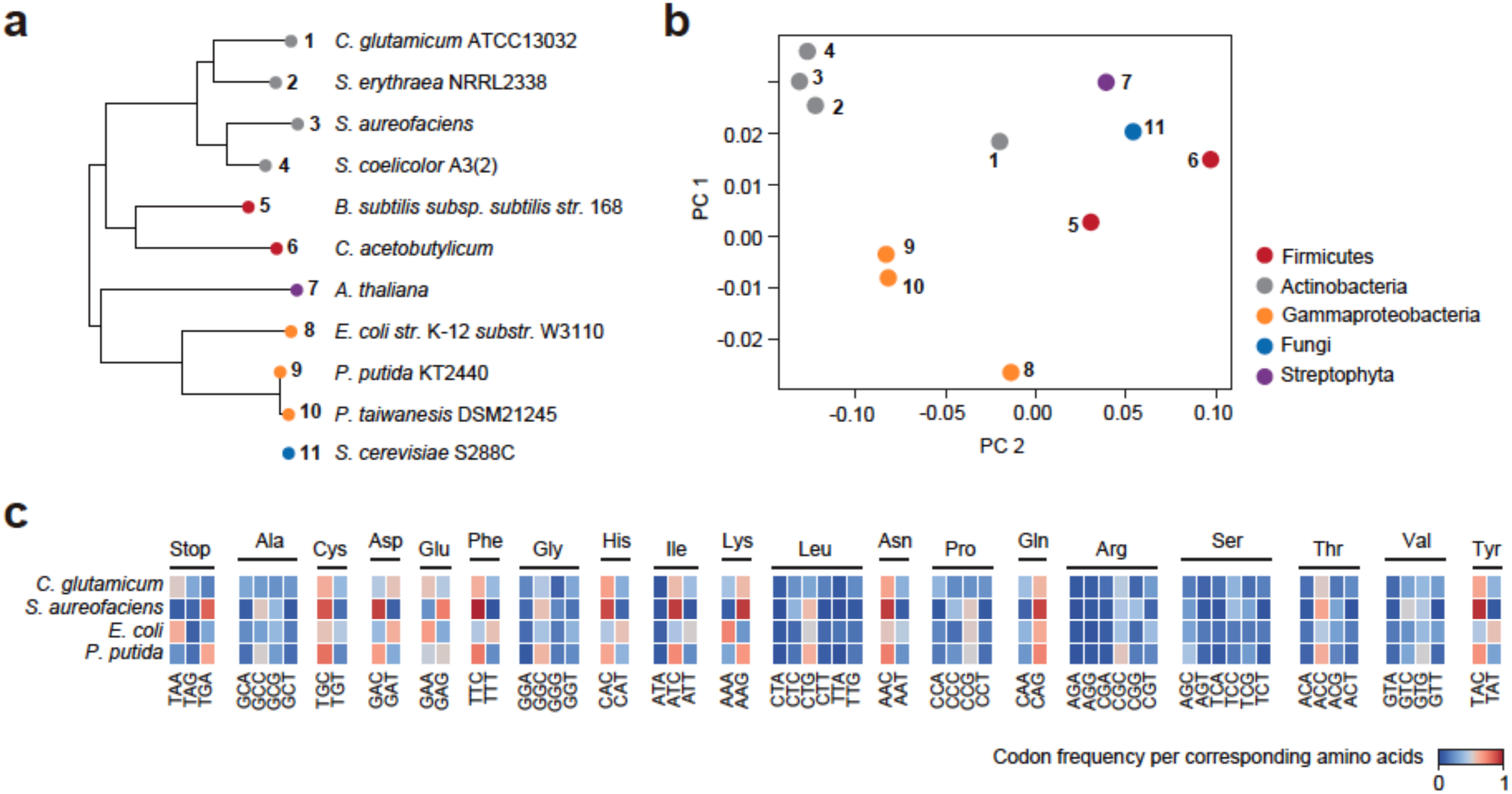
Global analysis of codon usage preferences between different species. (a) 16S rRNA sequence-based phylogenetic analysis of targeted hosts, native polyketide synthase hosts and outgroup references. *Saccharomyces cerevisiae* S288C was excluded. (b) Principal component analysis of codon usage tables associated with the same species from the phylogenetic analysis. (c) Comparison of the codon frequency per corresponding amino acid between *C. glutamicum*, *S. aureofaciens*, *E. coli*, and *P. putida*. The codons for methionine and tryptophan were excluded.

Principal component analysis (PCA) was performed on codon usage data for the same set of organisms (Figure 2b). The native PKS hosts *S. aureofaciens*, *S. erythraea*, and *S. coelicolor* are situated in close proximity to each other, whereas the evaluated heterologous hosts *C. glutamicum*, *E. coli*, and *P. putida* are more distant. Notably, despite *C. glutamicum* belonging to the same phylum, it does not cluster with the other actinobacteria. Furthermore, the significantly higher GC-content of *P. putida* (> 61 %) does not seem to lead to increased similarity in codon usage with streptomycetes either. Figure 2c highlights the relative codon frequency per corresponding amino acid (RCF) for *C. glutamicum*, *S. aureofaciens*, *E. coli*, and *P. putida*. While the RCF in *S. aureofaciens* has ten distinct maxima (> 0.8), *P. putida* is the only other organism with a similar maximum for the codon TGC. The high GC-content of *S. aureofaciens* (> 72 %) most likely leads to these maxima, especially for amino acids with only two codons. In contrast, the proteobacterium *E. coli* usually prefers codons with lower GC-content but seems to be more tolerable of other codons as well.

All in all, these data might suggest that there is no ideal host of industrial relevance that shows a similar codon preference with common native PKS hosts. PCA of codon usage could be a useful tool to assess the success rate of expression of the wildtype (WT) nucleotide sequence in the selected heterologous host. To further evaluate this theory, we performed codon optimization on an engineered PKS from *S. aureofaciens* Tü117 and compared expression levels with the WT nucleotide sequence in our selected heterologous hosts.

### Codon optimization of engineered lipomycin polyketide synthase

The design of our synthetic PKS was based on the work of Yuzawa et al. (2013). In short, we removed the first 59 N-terminal amino acids of the lipomycin PKS (LipPKS) and truncated module (M) 1 after the acyl carrier protein (ACP) 1. We then fused the remaining protein to the erythromycin PKS (EryPKS) M6 thioesterase (TE) including the interdomain linker between EryPKS ACP6 and TE6 (Figure 1). The N-terminal truncation leads to improved protein expression, and the TE hydrolyzes the product after the first methylmalonyl-CoA (mmCoA) extension (20). The loading domain of LipPKS is very promiscuous and can utilize isobutyryl-CoA, 2-methylbutyryl-CoA, isovaleryl-CoA and propionyl-CoA (19). The comparably small size and wide range of acceptable acyl-CoAs makes this engineered PKS a well-suited candidate for heterologous expression.

While this design has shown to yield a variety of short-chain 3-hydroxy acids *in vivo*, its functionality has not yet been evaluated in our chosen heterologous hosts. To increase the probability for functional expression of the engineered PKS, we employed the three codon optimization methods ‘use best codon’ (ubc), ‘match codon usage’ (mcu) and ‘harmonize relative codon adaptiveness’ (hrca) using DNA Chisel, and tested their effect on transcription and translation of the target PKS gene (15). A full list of the used settings for codon optimization can be found in the Supplementary (Table S1). Furthermore, we included the WT nucleotide sequence of each part of the PKS and investigated the effect of the two different start codons GTG and ATG.

As the applied algorithms are part of a Python-based toolkit with a command line interface (CLI), we developed a user-friendly graphical interface (GUI) to facilitate the use of open-source codon optimization tools. The codon optimization tool is publicly available, requires no prior experience in CLIs and can be reached under https://basebuddy.lbl.gov. In addition to the already implemented codon usage database from Kazusa (9), we also included the latest version of the Codon and Codon Pair Usage Tables (CoCoPUTs) database (21). Compared to the Kazusa database, CoCoPUTs are based on up-to-date sequencing data and also provide codon usage tables for a significantly broader range of organisms.

### 2.2 Development of a backbone excision-dependent expression system

To ensure greater accuracy and consistency of our results in *C. glutamicum* and *P. putida*, we utilized serine recombinase-assisted genome engineering (SAGE) to integrate our codon variants into the genome of these hosts (22, 23). The strains AG5577 and AG6212 are derived from *P. putida* and *C. glutamicum*, respectively, and contain a total of 9 unique *attB* sites in their genome (Adam Guss, personal communication). These genomic “landing pads” enable highly efficient and precise gene integrations via the SAGE system (Figure 3a).

**Figure 3:**
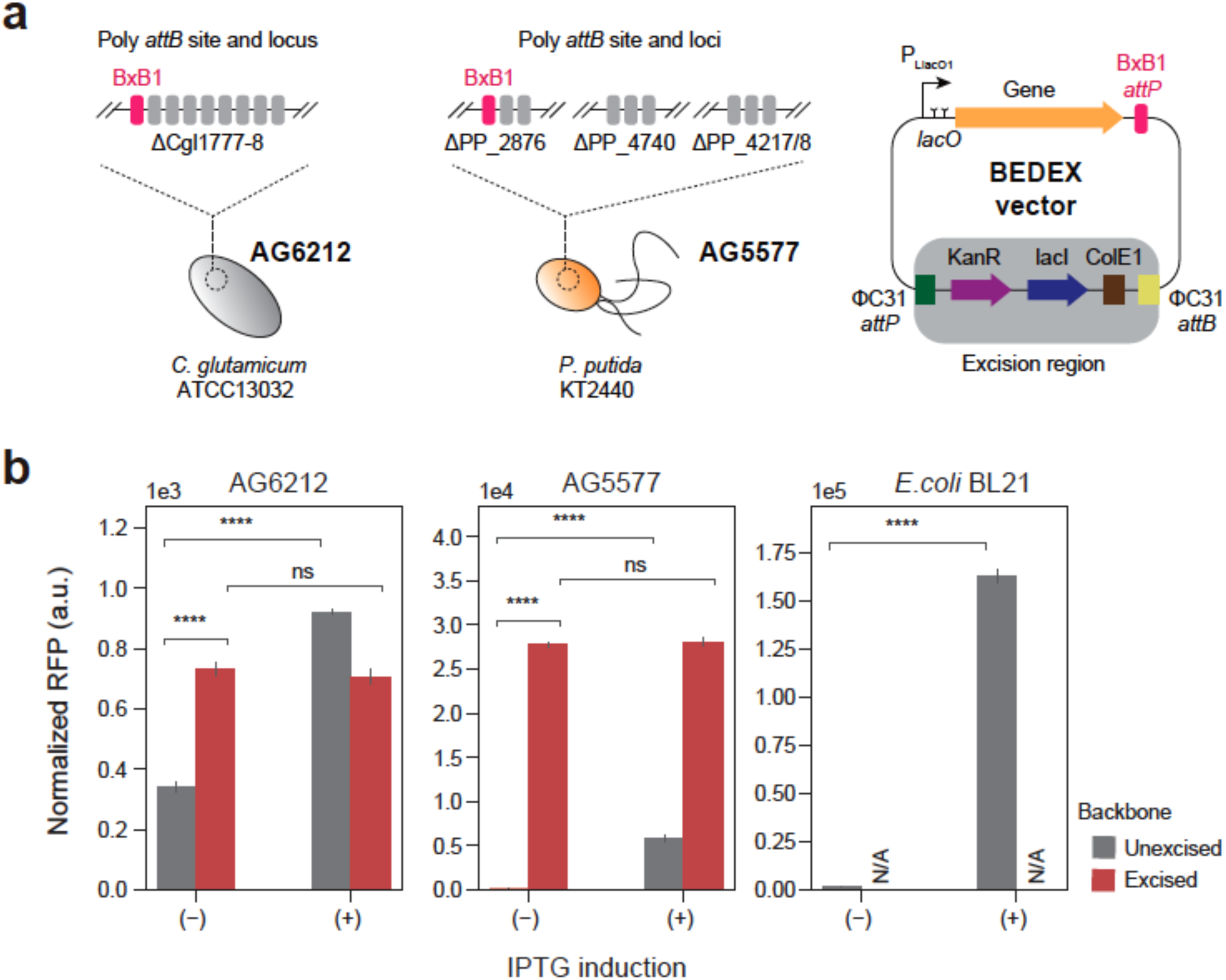
Extending the SAGE system with backbone excision-dependent expression (BEDEX) vectors. (a) *C. glutamicum* AG6212 and *P. putida* AG5577 contain a total of 9 unique *attB* sites each. Heterologous serine recombinases catalyze the integration of vectors containing the corresponding *attP* site. Expressing the integrase ΦC31 removes the integrated vector backbone and allows for selection marker recycling. Excising the backbone of BEDEX vectors also removes the repressor LacI. (b) *C. glutamicum* AG6212, *P. putida* AG5577, and *E. coli* BL21 containing BEDEX vector carrying RFP. Induction or excision of the BEDEX vector backbone leads to a significant increase in RFP levels (n = 3).

In addition, we sought to further improve the applicability of the SAGE system by combining it with backbone excision-dependent expression (BEDEX) vectors (Figure 3a). The two key elements of BEDEX vectors are the *LlacO1* promoter and the excisable *lacI* gene. In the case of the SAGE system, the integrated plasmid backbone is excised via transient expression of the ΦC31 integrase. By removing the *lacI* gene from the host genome, the *LlacO1* promoter is no longer repressed and becomes constitutive. To confirm the functionality of the BEDEX vectors in all three host organisms, we used red fluorescent protein (RFP) as a readily quantifiable output (Figure 3b). While the *rfp* gene was integrated into the genomes of *P. putida* AG5577 and *C. glutamicum* AG6212, we relied on plasmid-based expression in *E. coli* BL21. Due to the presence of the ColE1 origin of replication, the SAGE system vector suite cannot be utilized to engineer *E. coli*.

In *C. glutamicum* AG6212, the measured RFP signal was extremely low compared to *rfp*-expressing *E. coli* and *P. putida* cells. The *C. glutamicum* control sample showed a normalized RFP signal of 174 ± 4, which is about 2-fold lower than the uninduced and unexcised integration of the BEDEX vector (data not shown). However, induction or excision of the backbone showed a significant increase in RFP signal by up to 4-fold.

In *P. putida* AG5577, the expression was strongly repressed in the presence of *lacI*. The induction with 200 µM IPTG led to a 44-fold increase in normalized RFP signal. Excision of the backbone, however, led to a 213-fold increase in signal and the expression seemed unaffected by the addition of inducer. In *E. coli*, IPTG-induction achieved a 105-fold change in normalized RFP signal, while maintaining a relatively low signal of 1548 ± 287 in the uninduced state.

These data demonstrate the successful integration of the target gene into the genome of *C. glutamicum* and *P. putida*. In addition, by eliminating *lacI* from the plasmid backbone, we achieved expression without inducing the *LlacO1* promoter.

### 2.2 Quantification of heterologous protein and transcript

The BEDEX vectors carrying the 11 codon variants of LipPKS were introduced into *C. glutamicum* AG6212, *E. coli* BL21, and *P. putida* AG5577. The LipPKS gene was genomically expressed in *C. glutamicum* AG6212 and *P. putida* AG5577, and from plasmid DNA in *E. coli* BL21.

Proteomics analysis was performed after digesting the extracted proteins from cultured cells with trypsin. The final value for each protein represents the mean of the top 3 peptides or “best flyers” detected by LC-MS (24–27). To calculate the percentage of the total protein abundance, the intensity of the target protein was divided by the total sum of all detected protein intensities. The abundance of the engineered LipPKS for each codon version and host organism are shown in Figure 4a. For some *C. glutamicum* and *P. putida* constructs, it was not possible to excise the genomically integrated backbone of the BEDEX vector. The data for these strains has been highlighted by hatching the corresponding bar plot (Figure 4a).

**Figure 4:**
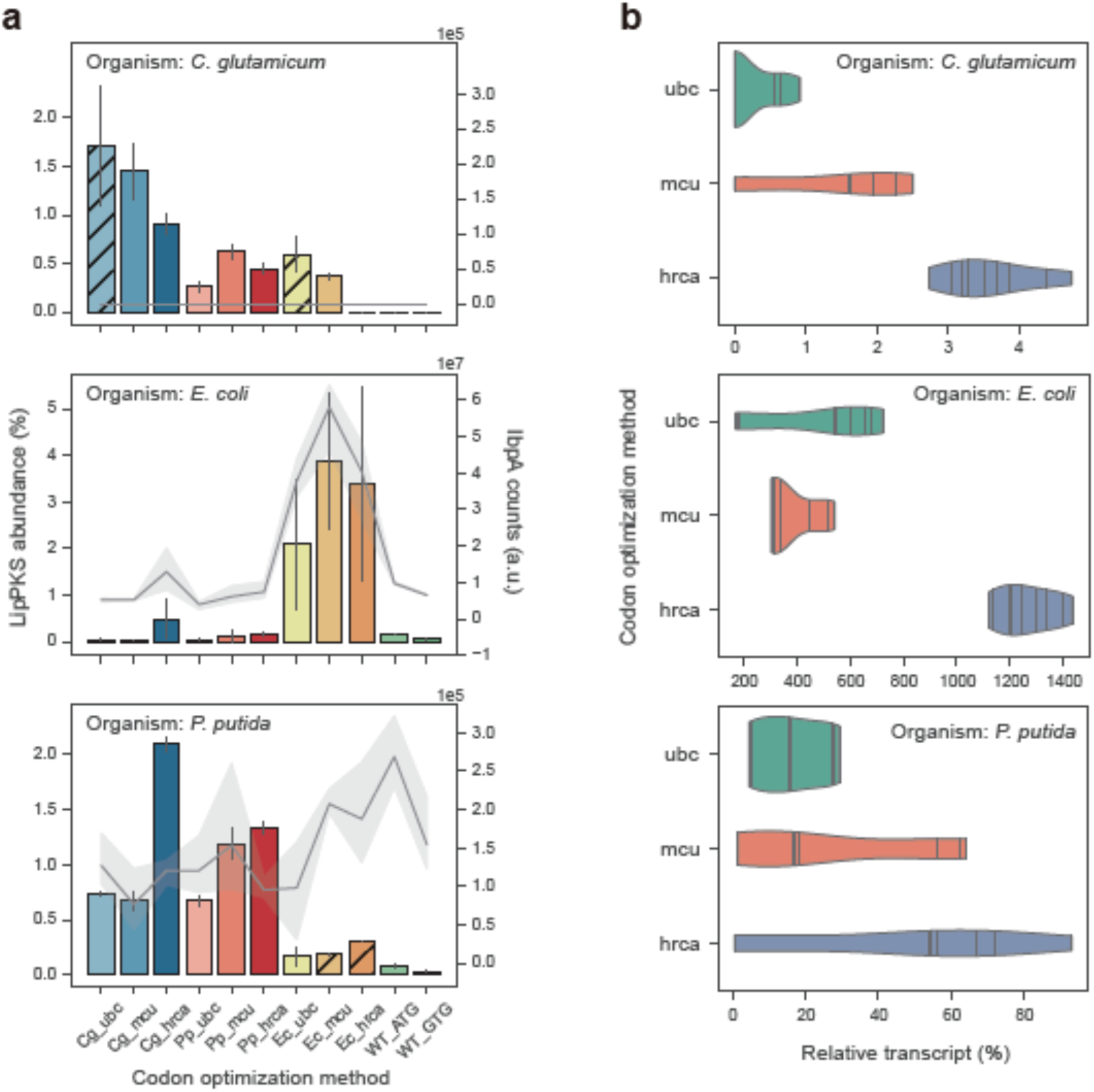
Relative abundance of LipPKS peptides and transcript in heterologous hosts expressing different codon variants of LipPKS. (a) Calculated protein abundance (n = 3) is the relative intensity of the top 3 peptides that correspond to the target protein divided by the intensity of all proteins detected. Hatched bars indicate strains with non-excisable backbones. Levels of the insolubility marker IbpA are represented by a gray line (n = 3). IbpA is not present in *C. glutamicum*. (b) Each violin represents the distribution of the relative transcript amount for all target host optimizations with the same optimization strategy (n = 9). Relative transcript was calculated from the Ct value difference between target transcript and housekeeping gene. The lines within the violins are individual data points. Target host optimization: Cg = *C. glutamicum*; Ec = *E. coli*; Pp = *P. putida*; Optimization strategy: ubc = use best codon; mcu = match codon usage; hrca = harmonize relative codon adaptiveness

The LipPKS counts in *C. glutamicum* samples were on average 1 to 2 orders of magnitudes lower than in *E. coli* and *P. putida* samples. However, if we compare protein abundance instead, values become more similar between organisms. Depending on the host and codon version, LipPKS abundance ranges from 0 to 3.9 ± 1.5 %. In general, PKS genes codon optimized for a particular organism produced the most PKS when expressed in the target organism. For example, PKS genes that were codon optimized for *P. putida* (Pp) and were expressed in *P. putida* achieved a LipPKS abundance of 1.1 ± 0.4 % across all three codon optimization methods. When compared to PKS genes that were codon optimized for *E. coli* (Ec) but expressed in *P. putida*, LipPKS abundance was as low as 0.2 ± 0.1 %. This trend was even more drastic when PKS genes codon optimized for all three organisms were expressed in *E. coli*: the *E. coli* codon optimized genes produced on average 22-fold more LipPKS than the genes codon optimized for *P. putida* and *C. glutamicum*.

Another noticeable difference was the relative standard deviation between plasmid-based and genomic expression, which can be as high as 121 % or as low as 2%, respectively. These data suggest that the expression of LipPKS from plasmid DNA can lead to greater data variability and might be less reproducible between replicates.

In general, the WT nucleotide sequences performed very poorly: protein levels were either undetectable or extremely low compared to the codon optimized variants. However, mutating the original GTG start codon to ATG resulted on average in a 3-fold increase in peptide counts.

In addition to the LipPKS abundance, we also determined the IbpA levels for *E. coli* and *P. putida* samples (Figure 4a). The small heat-shock protein IbpA has been shown to be expressed in the presence of insoluble protein in the cell. Generally, a high IbpA level is an indicator for misfolded protein and can give a hint about the activity of the heterologously produced protein in these hosts (28). In *E. coli*, there is a strong Pearson correlation of 0.94 between IbpA and LipPKS levels, whereas no such correlation was observed in *P. putida*. Interestingly, expressing the WT_ATG construct in *P. putida* resulted in the highest IbpA signal, while producing a relatively low amount of LipPKS (0.09 ± 0.02 %). Therefore, the poor similarity in codon usage may have resulted in not only low levels of protein but also insoluble protein.

To date, it is not clear if codon usage affects gene expression on a transcriptional or translational level, especially across different organisms (29, 30). To analyze the correlation between transcript and target protein, we measured the LipPKS transcript level by reverse transcription qualitative real-time PCR (RT-qPCR). Transcript levels were calculated from the Ct value difference between target transcript and housekeeping gene (Supplementary Table S2). The strongest correlation between transcript and protein levels was observed in *P. putida* (R = 0.67), while the correlation coefficient for the other two hosts was below |0.3| (Supplementary Figure S1). In general, a low transcript amount led to a low protein level, with the exception of the codon variants Cg_ubc, Cg_mcu, and Ec_ubc in *C. glutamicum*. Here, we were not able to detect any transcript, although we detected significant amounts of LipPKS peptides. This was also the case for Ec_mcu and Ec_hrca variants in *P. putida*. With the exception of the Cg_mcu variant in *C. glutamicum*, all these variants were induced at the time of inoculation, which might have affected mRNA levels.

Another noticeable trend was the high transcript amount for the hrca codon optimization, which seemed to be independent of the targeted host (Figure 4b). This trend was especially significant in *E. coli* where hrca codon optimizations yielded 1266 ± 111% relative transcript, while ubc optimizations resulted in 378 ± 96 % relative transcript.

Interestingly, for the codon variants Pp_hrca and Cg_hrca, the high transcript amount did not lead to high protein levels. These findings support the hypothesis that codon frequency might determine translation rates, as evidenced by the higher LipPKS counts of the target host variants of hrca optimizations.

### 2.3 Production of an unnatural polyketide by engineered PKS

The production of the PKS protein itself does not require any genetic modifications of the chosen hosts. However, to demonstrate that our engineered PKS is functional and produces the desired polyketide, we needed to express supplementary pathways. For successful polyketide production in *E. coli*, the PKS vectors were transformed into the readily available host *E. coli* K207-3 (31). This strain possesses the required mmCoA pathway and phosphopantetheinyl transferase (PPTase) for the supply of the extender substrate and activation of the ACP domain. Due to the versatile nature of the LipPKS loading module, it exhibits promiscuity towards propionyl-CoA as an alternative starter unit, making the expression of an isobutyryl-CoA pathway unnecessary (19).

Given *P. putida*’s demonstrated broad specificity PPTase and its ability to utilize the isobutyryl-CoA precursor valine, the integration of a heterologous mmCoA pathway is the sole modification required (32, 33). As previously reported, the mmCoA mutase and epimerase from *Sorangium cellulosum* So ce56 is functional in *P. putida* (*34*). Therefore, we integrated the unmodified operon under the control of the *tac* promoter into the MR11 *attB* site of *P. putida* AG5577. Unlike the BEDEX vector, the selected vector backbone lacks *lacI*, which eliminates the need for backbone excision (Figure 5a).

**Figure 5:**
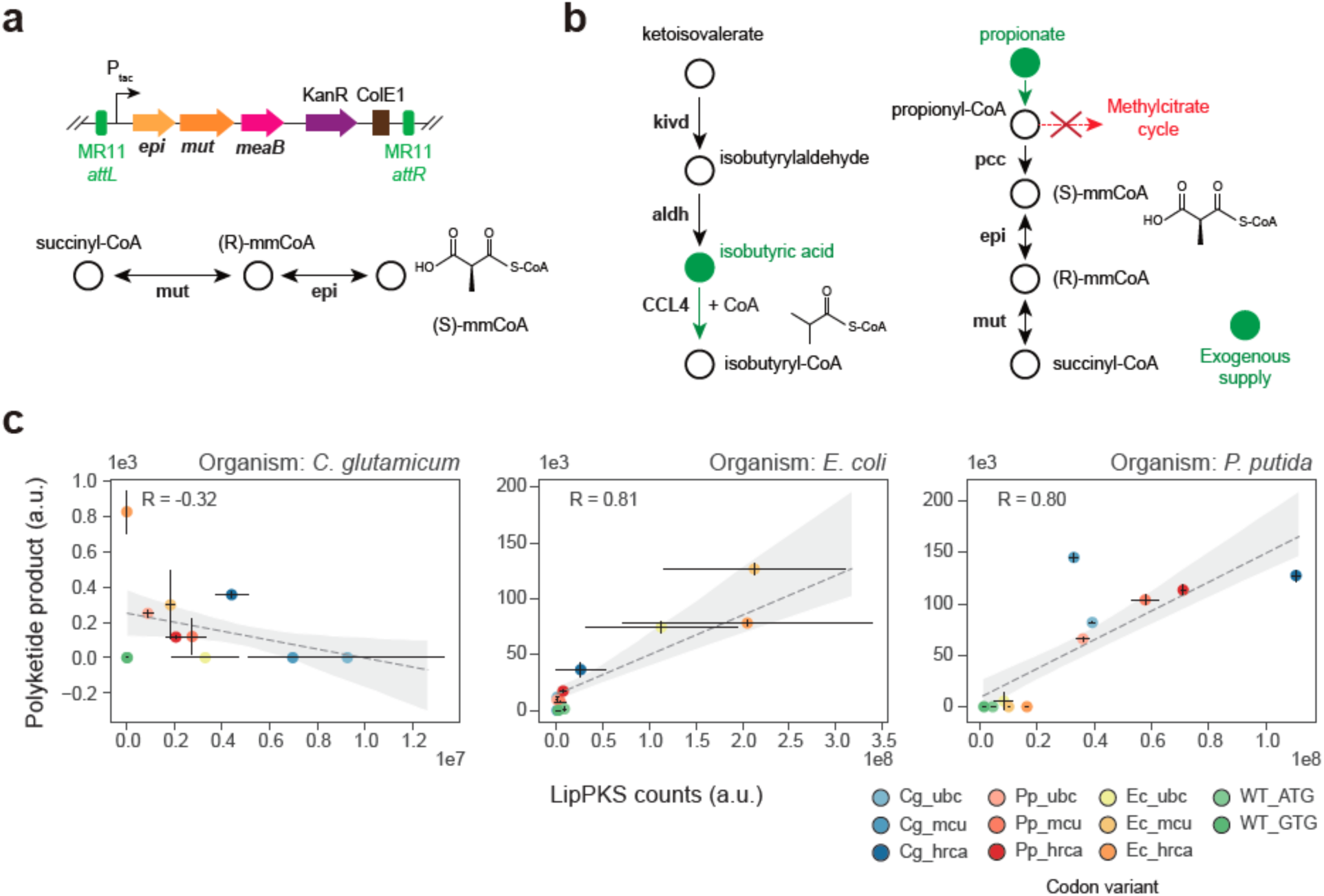
Production of an unnatural polyketide by engineered heterologous hosts. (a) Serine recombinase-assisted integration of the methylmalonyl-CoA mutase (mut) and epimerase (epi) pathway into *P. putida* AG5577. mmCoA: methylmalonyl-CoA (b) Engineered pathway for the production of the LipPKS loading substrate isobutyryl-CoA by the heterologous enzymes Kivd (ketoisovalerate decarboxylase) and CCL4 (2-methylpropanoate--CoA ligase) in *C. glutamicum* AG6212. Isobutyric and propionic acid were added to the *C. glutamicum* production medium (green). aldh: endogenous aldehyde dehydrogenase activity. (c) Regression plot of the LipPKS counts and polyketide production levels in the engineered hosts *C. glutamicum* AG6212cz, *E. coli* K207-3, and *P. putida* AG5577mm. Target host optimization: Cg = *C. glutamicum*; Ec = *E. coli*; Pp = *P. putida*; Optimization strategy: ubc = use best codon; mcu = match codon usage; hrca = harmonize relative codon adaptiveness

The third host organism, *C. glutamicum*, cannot utilize valine as the sole source of carbon, and the existence of an isobutyryl-CoA pathway is unlikely. However, previous studies have confirmed the presence of mmCoA, while the presence of propionyl-CoA is highly probable (35). Unfortunately, it remains unclear if there is a type I PKS compatible PPTase in *C. glutamicum*. To address this, we employed conventional cloning techniques to introduce the PPTase *sfp* into *C. glutamicum* AG6212. Furthermore, to increase the probability of producing any kind of polyketide, we also integrated the heterologous CoA ligase *CCL4* and the ketoisovalerate decarboxylase *kivd*. By relying on endogenous aldehyde dehydrogenase activity, both genes combined could potentially lead to the preferred LipPKS starter unit isobutyryl-CoA (36–38). In addition, we added isobutyric and propionic acid to the production medium to increase precursor availability. Finally, the gene *prpDBC2* was removed to facilitate flux through mmCoA by disrupting the methyl citrate cycle (Figure 5b).

The LipPKS protein level and polyketide production was measured in all three hosts (Figure 5c). The production of the corresponding polyketide was confirmed by an authentic standard (Supplementary Figure S2). In *C. glutamicum* and *P. putida*, we chose to focus on the production of (3R,2R)-3-hydroxy-2,4-dimethylpentanoic acid (3H24DMPA), although, in *P. putida*, we identified the corresponding masses for the polyketides made with the starter units 2-methylbutyryl-CoA and propionyl-CoA as well (Supplementary Figure S2a). In *E. coli*, the only possible product is (3R,2R)-3-hydroxy-2-methylpentanoic acid (3H2MPA).

In *P. putida* and *E. coli*, the protein and product levels of the codon variants showed a Pearson correlation coefficient of 0.80 and 0.81, respectively. The highest product titer for *P. putida* was achieved by the Cg_mcu codon variant, which was a noticeable outlier when compared to other protein and product relations. In *E. coli*, the highest product titer was achieved by the Ec_mcu codon variant. Interestingly, our *in vivo* results in *P. putida* confirm the malonyl-CoA extended product (Supplementary Figure S2), which contradicts the reported and predicted mmCoA specificity of LipPKS AT1 (19).

Polyketide production in *C. glutamicum* was very low and barely detectable (Supplementary Figure S2b), despite the significant LipPKS levels. There was also no clear correlation between LipPKS peptides and polyketide product (R = -0.32). In addition to analyzing LipPKS amounts in this host, we also measured the peptide counts for Sfp, Kivd and CCL4. However, we were only able to detect peptides that correspond to Sfp and Kivd; CCL4 peptides were not detected (Supplementary Figure S3). Therefore, the strain might be missing the CoA ligase to efficiently activate the starter unit isobutyric acid.

## 3 DISCUSSION

In this study, we identified the most suitable codon optimization method for an engineered T1PKS for heterologous expression in *C. glutamicum*, *E. coli*, and *P. putida*. Furthermore, we elucidated the relationship between codon variant, protein level, transcript amount and product titer.

The Python library DNA Chisel allowed us to do fully customizable and transparent codon optimizations of our target gene. The degree of customization was an important factor to find an appropriate balance between optimization task and synthesis difficulty. For instance, the implementation of the UniquifyAllKmers constraint can greatly reduce the amount of repetitive sequence, which remains a major hurdle in the chemical synthesis of DNA (39). As a result, codon optimization efficiency may be constrained by the limitations of current DNA synthesis techniques.

While the Kazusa codon usage database was sufficient in our case, it might become a limiting factor for future investigations of more exotic PKSs or heterologous hosts. The ubc and mcu methods only require the targeted host’s codon usage, whereas the hrca method also requires a codon usage table of the source organism. However, our developed online tool includes the database CoCoPUTs, which greatly improves the codon usage table accuracy and the number of available organisms. During codon optimization, it is often unclear which codon usage database is used, especially by commercialized codon optimization tools. This can make a significant difference in the accuracy of the optimized gene. For example, the *S. aureofaciens* (NCBI:txid1894) codon table from Kazusa contains only 80 CDSs, while the codon table for the same organism on CoCoPUTs contains 37,337 CDSs. The substantial discrepancy is most likely a result of recent advances in genome sequencing and, therefore, more accurate bacterial genotyping (40).

Given that our systematic approach amounted to 11 codon variants of a relatively large gene (>7 kb), we sought to avoid relying exclusively on plasmid-based expression systems. Besides well-known issues such as plasmid stability, intercompatibility between our different organisms might have become challenging as well. The SAGE system has proven to be an indispensable tool for genetic engineering when reproducibility and throughput are limiting factors. Stable integrations of large genes into the chosen host genomes can be multiplexed, are highly efficient and require minimal knowledge about the targeted host. In combination with BEDEX vectors this system also enables precise control over gene expression and can improve plasmid assembly efficiency by mitigating the cellular burden of strong constitutive promoters or potentially toxic gene products (41). We have also shown that BEDEX vectors are universally functional in our chosen hosts, although measured RFP levels in *C. glutamicum* were very low. This observation could have multiple reasons, including a thicker cell wall preventing the emission of fluorescence or simply a non-compatible ribosome binding site (RBS) or promoter (42). Nonetheless, the measured differences between the excised and unexcised vector backbone were significant, and PLlacO1 became constitutively active in this host.

In an effort to further evaluate the BEDEX system, we applied it to construct our LipPKS expression strains. As confirmed with RFP, the BEDEX system showed a similar performance during the construction process. However, we encountered some difficulties while excising the backbone of certain codon variants. The presence of highly incompatible codon variants could result in growth inhibition, which then leads to the selection for *lacI* positive strains. In an attempt to investigate this behavior, we expressed the codon variants Pp_mcu and Ec_mcu from an inducible vector system in the host *P. putida.* As previously shown, the Pp_mcu codon optimized variant produced a significant amount of LipPKS peptides and product, whereas the Ec_mcu variant only produced the associated peptides (Figure 4a and 5c). During plasmid-based expression of these LipPKS variants, the protein levels for each variant were comparable. However, the levels of the insolubility marker IbpA were significantly higher in *P. putida* expressing the Ec_mcu LipPKS than in the strain expressing the Pp_mcu LipPKS (Supplementary Figure S4). Furthermore, we were not able to detect any product formation with the Ec_mcu codon variant. This could be an indicator for the expression of insoluble or misfolded protein by an incompatible codon variant.

The measured differences in LipPKS protein amounts clearly show that codon usage bias is a crucial factor in heterologous expression of PKSs. Compared to the WT codon version, our best performing codon variants exhibited a minimum 50-fold increase in PKS protein levels, which was also essential to enable the detection of the corresponding polyketide. However, we do not fully understand the cause of these differences. In *P. putida*, our RT-qPCR results might suggest that there is a link between codon usage and transcript level, whereas the *E. coli* and *C. glutamicum* data show no such correlation. In the two later hosts, codon frequency might have a stronger influence on the protein level. Current literature presents evidence for both of these theories (29, 30). While our investigation could not provide a satisfying answer to this controversial topic, we still identified an optimal way for codon optimization of PKSs. According to our data, the host-specific mcu method resulted in the highest protein and product levels, whereas the hrca method gave rise to the highest transcript levels. The choice of the most optimal algorithm is therefore dependent on the goal or suspected limitation of the host. The difference in protein production using ATG and GTG start codons could also be confirmed, and our results are consistent with prior studies (43). Utilization of GTG start codons seems to be more prevalent in high GC organisms and the percentage of GTG initiation codons decreases to less than 25 % in lower GC organisms (<65 %) (44). For heterologous expression of PKSs, the G to A mutation can be easily achieved by site-directed mutagenesis and is a cost-effective way to improve expression levels.

In addition to *E. coli*, the two other tested hosts have proven to be suitable for the expression of engineered T1PKSs. The activity of the PKS and the production of the expected 3-hydroxy acid product could be confirmed in all three hosts. Since we detected sufficient levels of protein in *C. glutamicum*, it can be assumed that the exceedingly low product levels are a result of the insufficient supply of isobutyryl-CoA. The loading domain of LipPKS has a strong preference for isobutyryl-CoA but does also accept 2-methylbutyryl-CoA, isovaleryl-CoA, and propionyl-CoA (19). However, the catalytic efficiency of utilizing propionyl-CoA is about 4-fold lower than compared to isobutyryl-CoA. It is also unclear whether *C. glutamicum* metabolizes short-chain 3-hydroxy acids, which could be another reason for the low product titer. In addition, the employed engineering strategy needs further evaluation of intracellular acyl-CoA concentrations and precursor pathways.

While we successfully identified the most effective strategy for codon optimization of a T1PKS, we still lack a complete understanding of the underlying reasons for its success. Such understanding may only come by codon optimizing many different types of genes in many different hosts, a project that would be economically infeasible with current DNA synthesis costs. In addition, it is possible that there are even better methods for codon optimizing genes; indeed, recent developments in artificial intelligence and machine learning could play a crucial role in optimizing heterologous gene sequences in the future (45).

## 4 METHODS

### 4.1 Chemicals, media and culture conditions

All chemicals were purchased from Sigma-Aldrich (USA) unless otherwise described. The authentic standards for (3R,2R)-3-hydroxy-2,4-dimethylpentanoic acid, 3-hydroxy-4-methylpentanoic acid, (3R,2R)-3-hydroxy-2-methylpentanoic acid, and 3-hydroxy-2,4-dimethylhexanoic acid were synthesized by Enamine (Ukraine).

Precultures of *E. coli*, *C. glutamicum* and *P. putida* were inoculated from a single colony and grown overnight. For *P. putida* and *E. coli* cultures, we used lysogeny broth (LB) medium and temperatures of 30 and 37 °C, respectively. For *C. glutamicum* cultures, we used brain heart infusion (BHI) medium and a temperature of 30 °C. If applicable, 50 μg/mL of kanamycin was added to the medium. The conditions for precultures and main cultures were kept consistent throughout this study unless otherwise described. For main cultures, 2 mL LB or BHI medium was inoculated with 20 µL preculture and grown in a 24-well plate (VWR, USA). Furthermore, no antibiotic was added to cultures of strains that contained a genomically integrated selection marker. The shaking speed was set to 200 rpm. The length and temperature of the cultivation were dependent on the organism and the sample requirements of the analyses. Strains that were containing *lacI* were induced by adding 200 µM IPTG to the medium.

### 4.2 Plasmids and strains

Plasmids and strains used in this study can be found in Table 1. All strains and plasmids generated in this work are publicly available through the JBEI registry (https://public-registry.jbei.org/folders/789). The codon optimized gene sequences of LipM1 fused to EryM6-TE were synthesized and cloned into the pBH026 vector backbone by Genscript (USA). The WT nucleotide sequences of LipM1 and EryM6-TE were obtained from *S. aureofaciens* Tü117 and *S. erythraea* NRRL2338 genomic DNA, respectively. Gibson primers were designed using the j5 software (46). PCR products and NdeI/XhoI (NEB, USA) digested pAN001 vector DNA were assembled using Gibson assembly standard protocol. Assembly of pGingerBG-NahR constructs was conducted with PCR amplified vector DNA. Plasmids were isolated with the Qiaprep Spin Miniprep kit (Qiagen, Germany). Primers were synthesized and purchased from IDT (USA). A list of all used primers can be found in the Supplementary (Table S2).

**Table 1:**
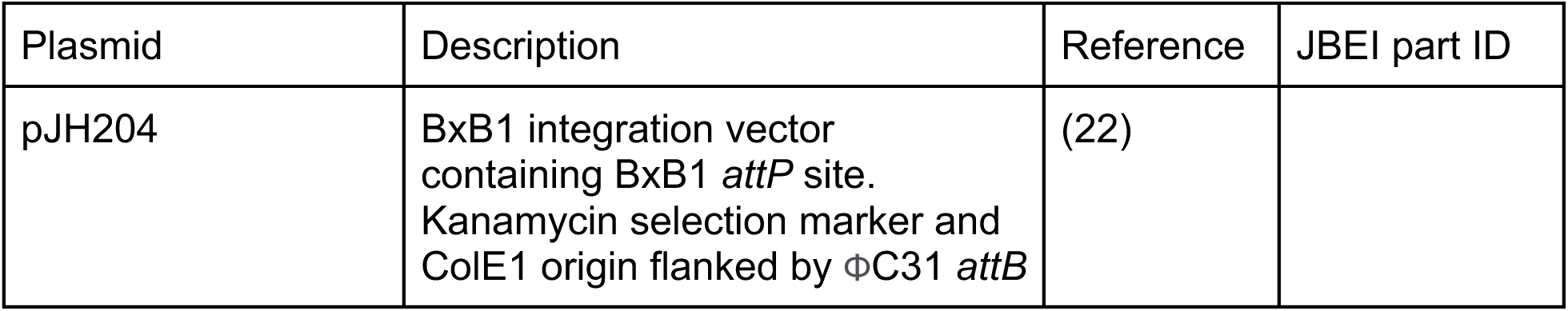
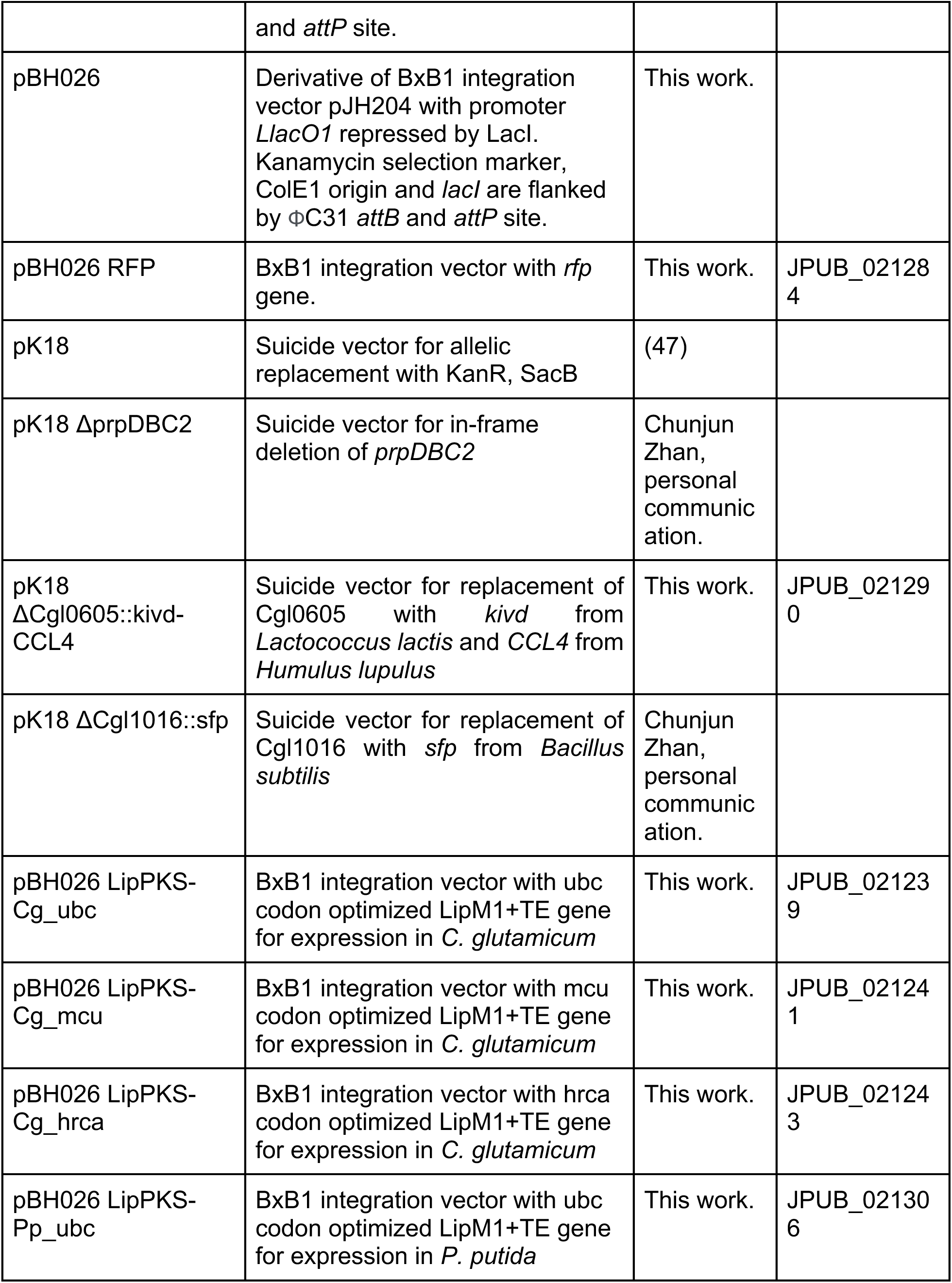
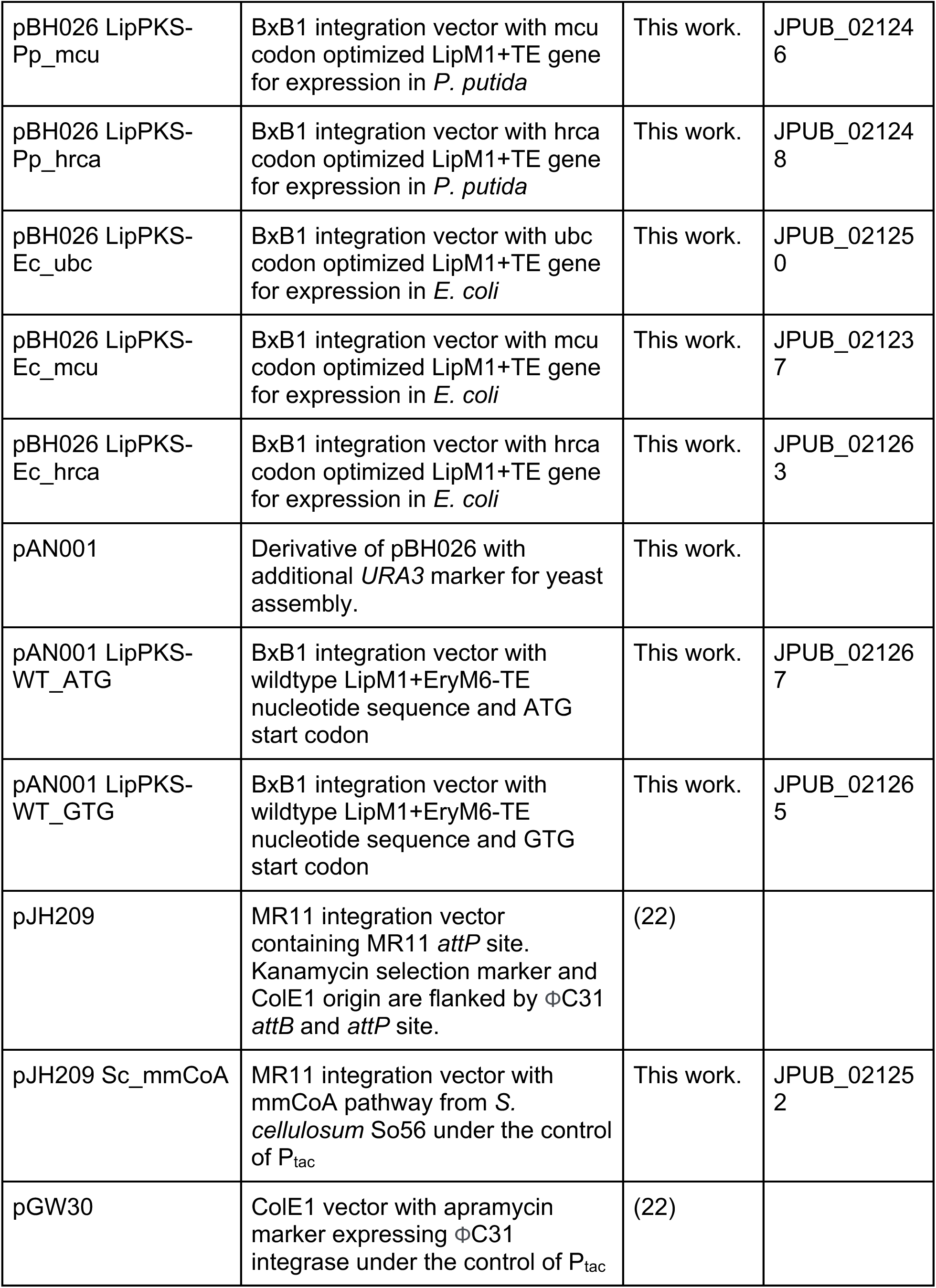
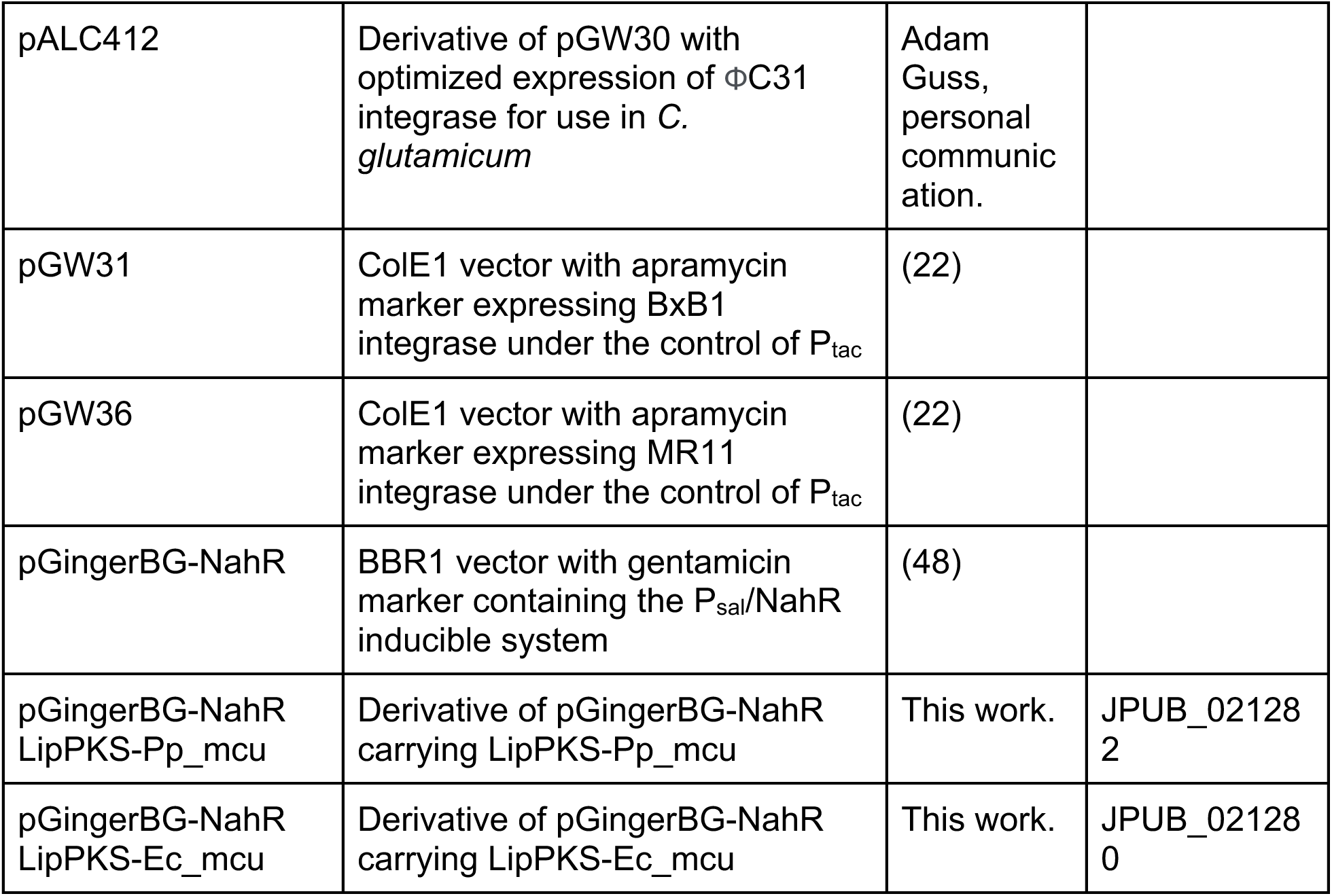
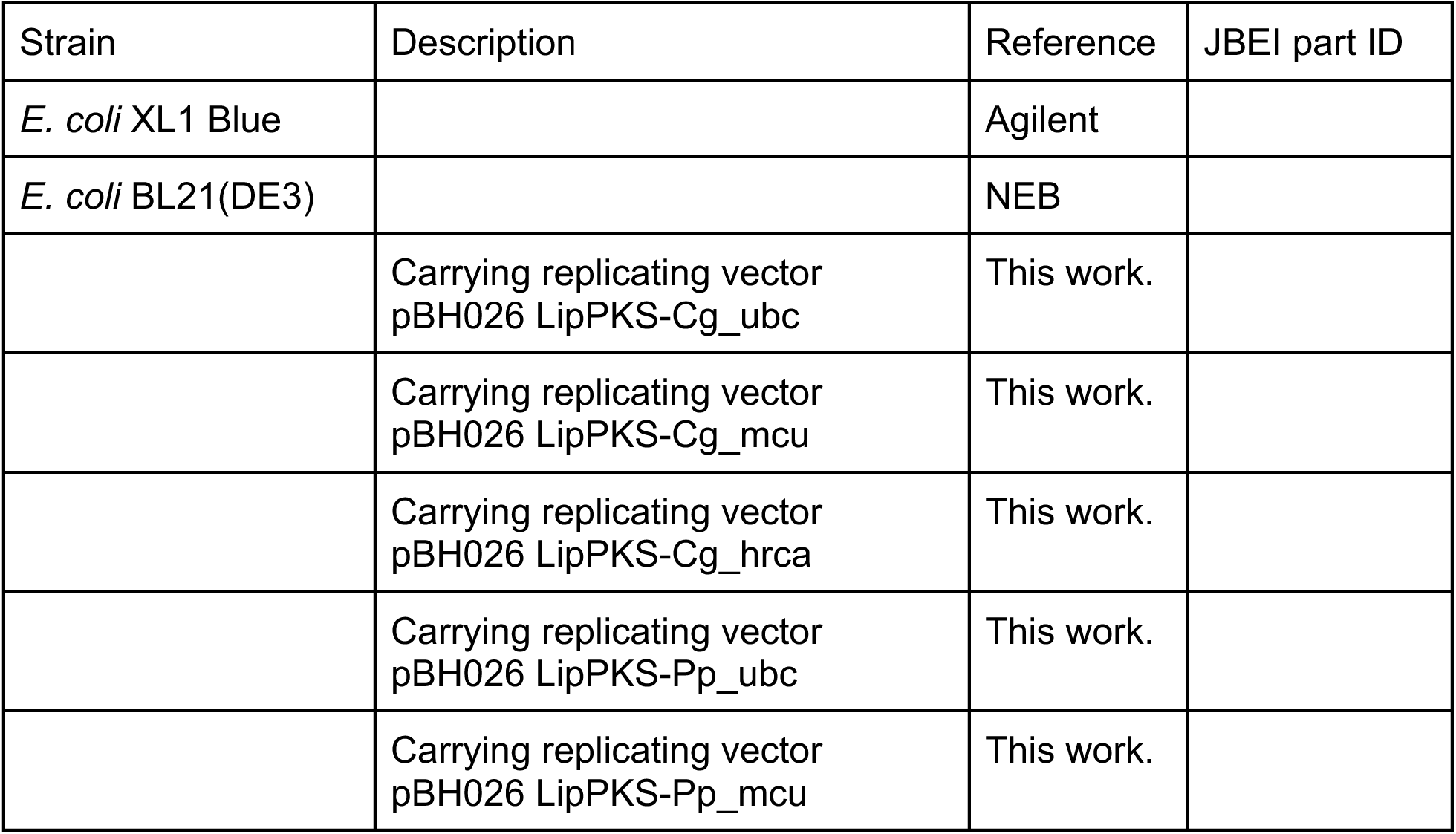
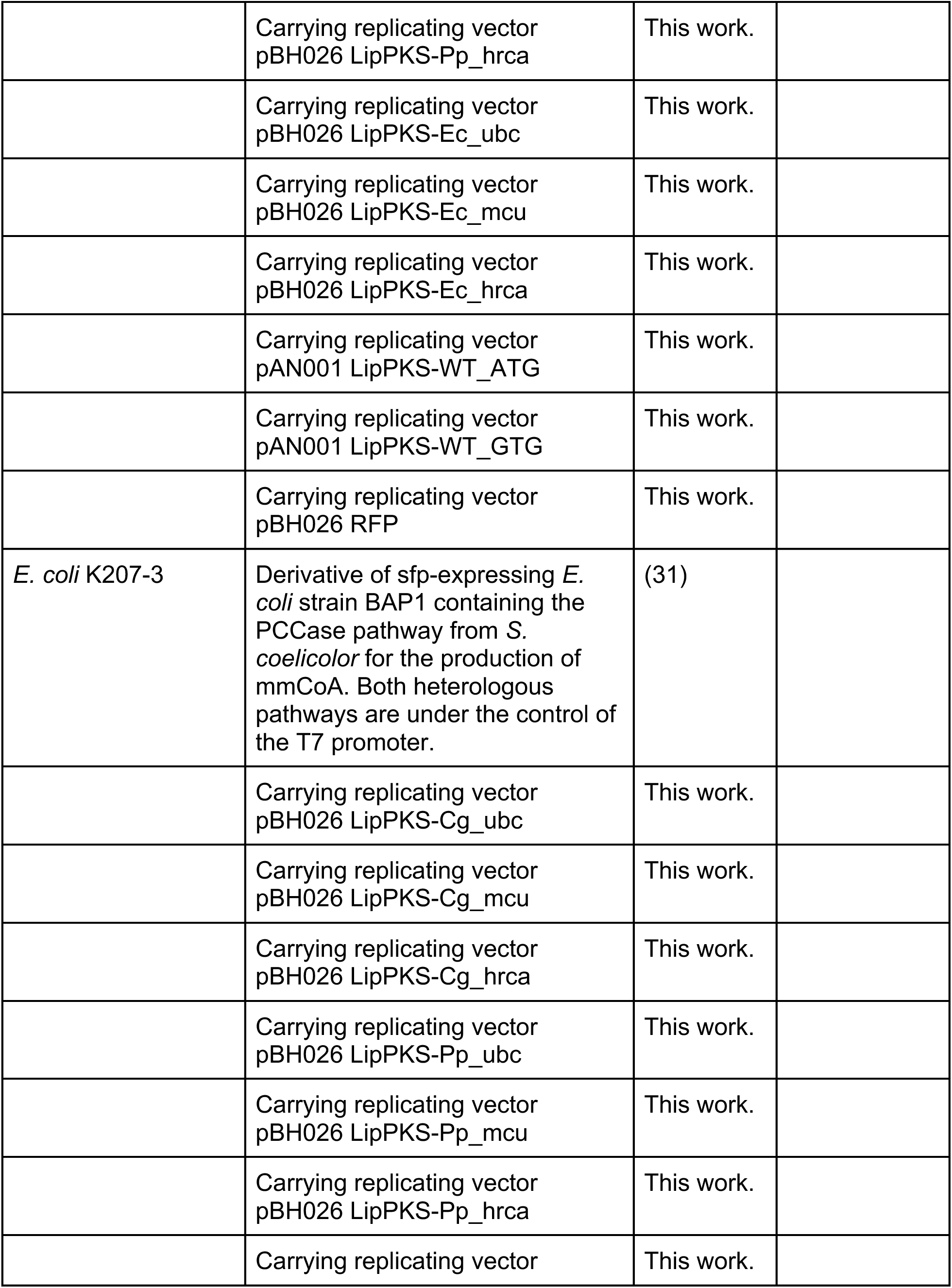
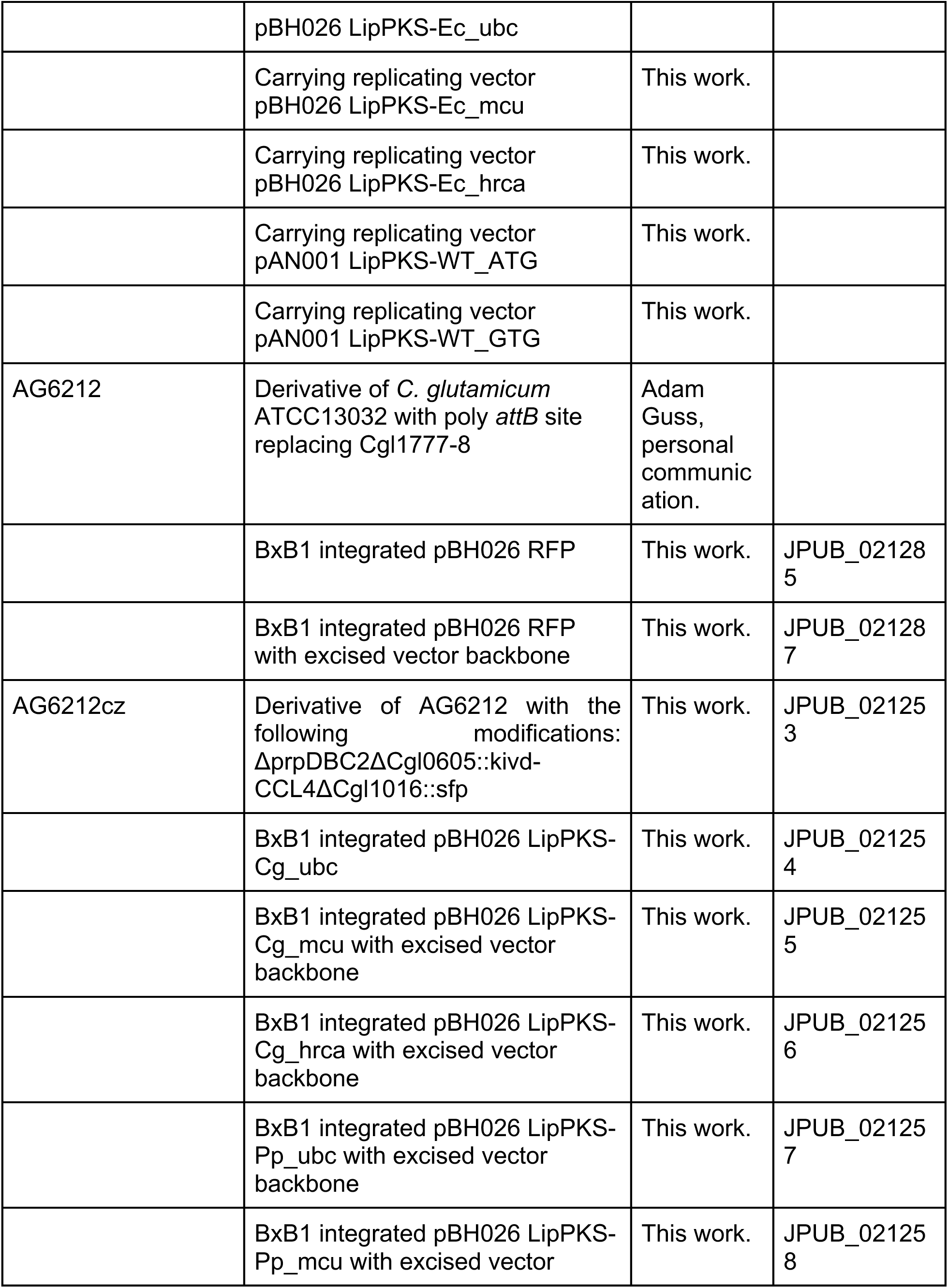
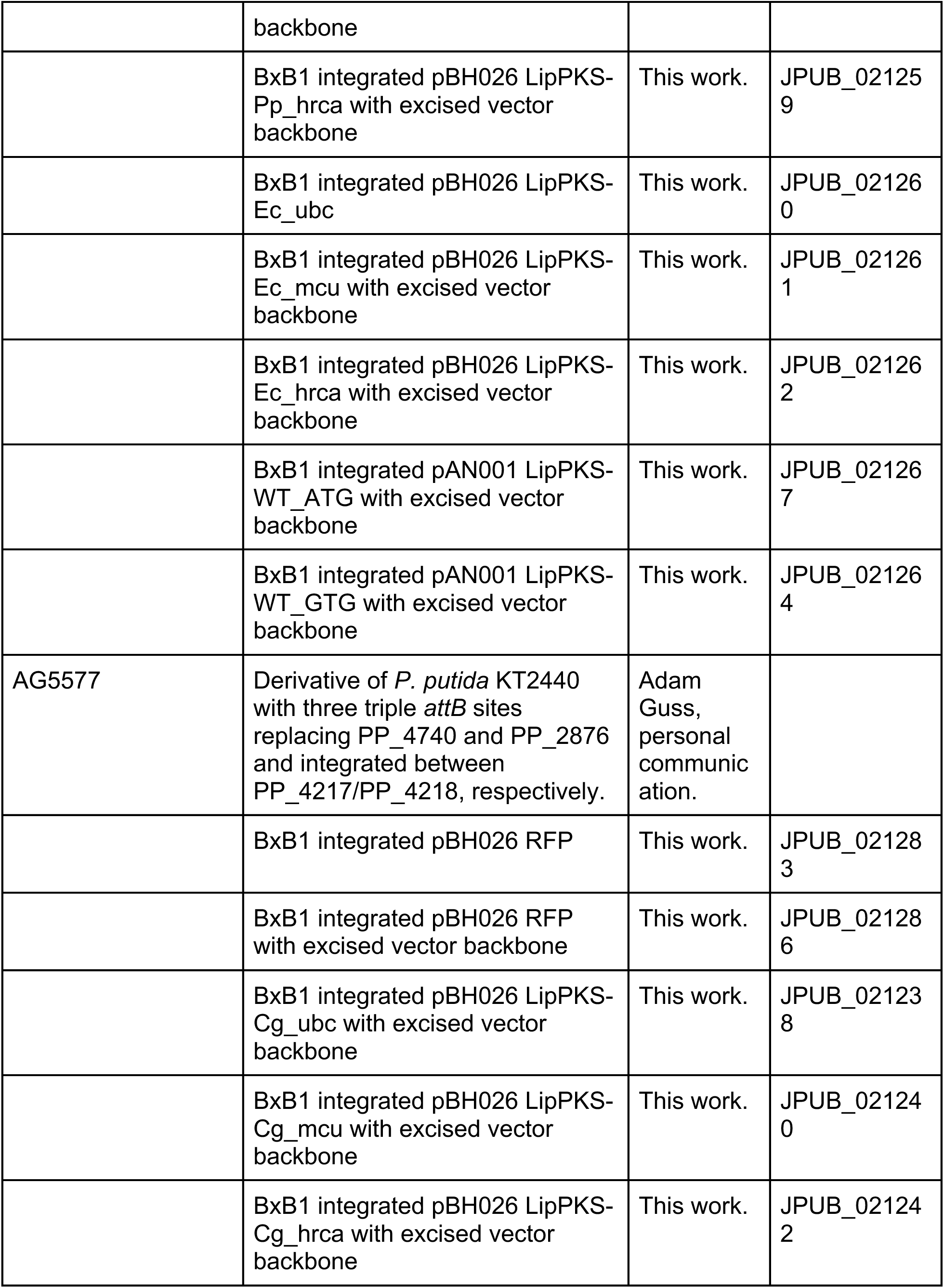
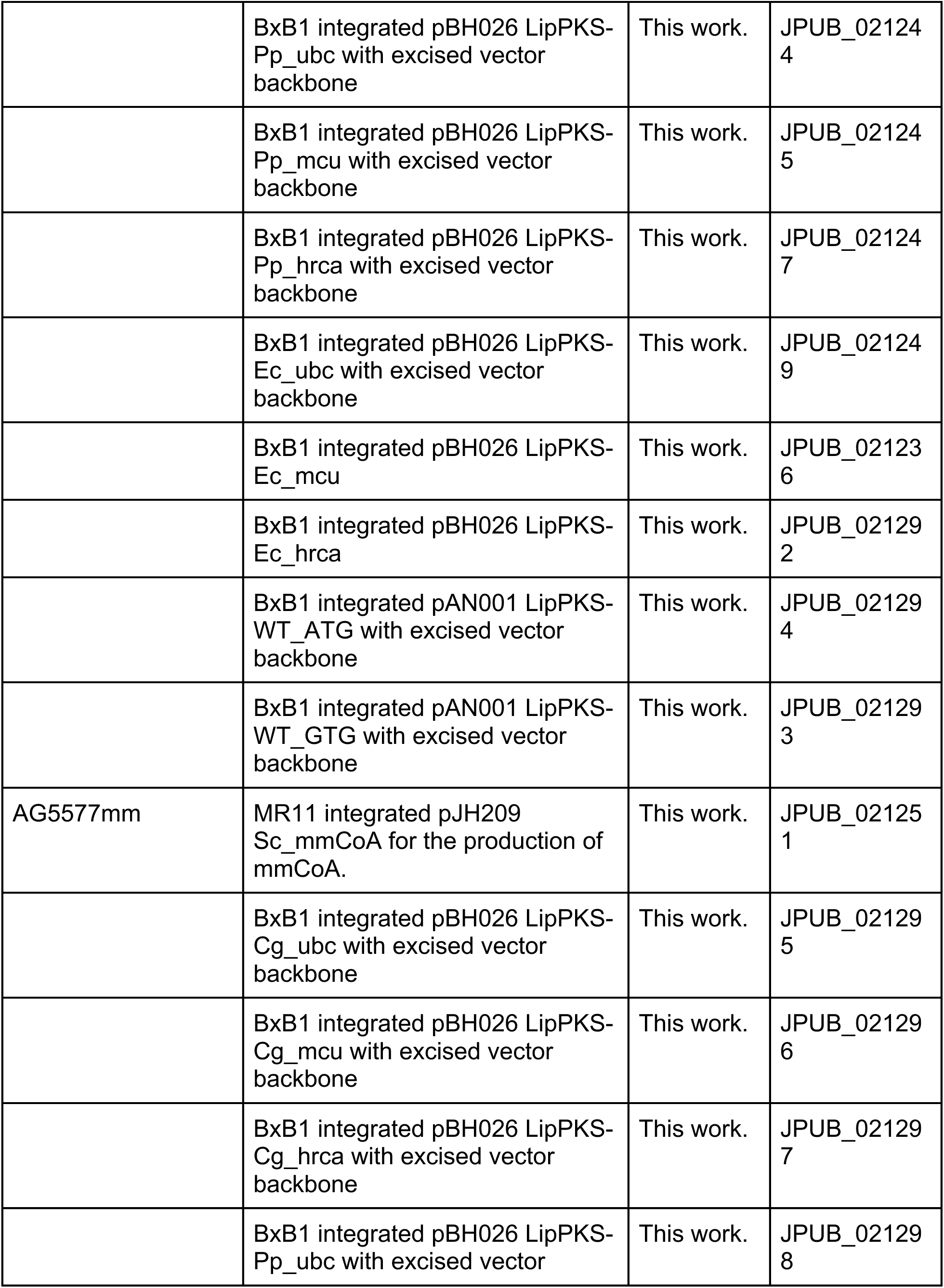
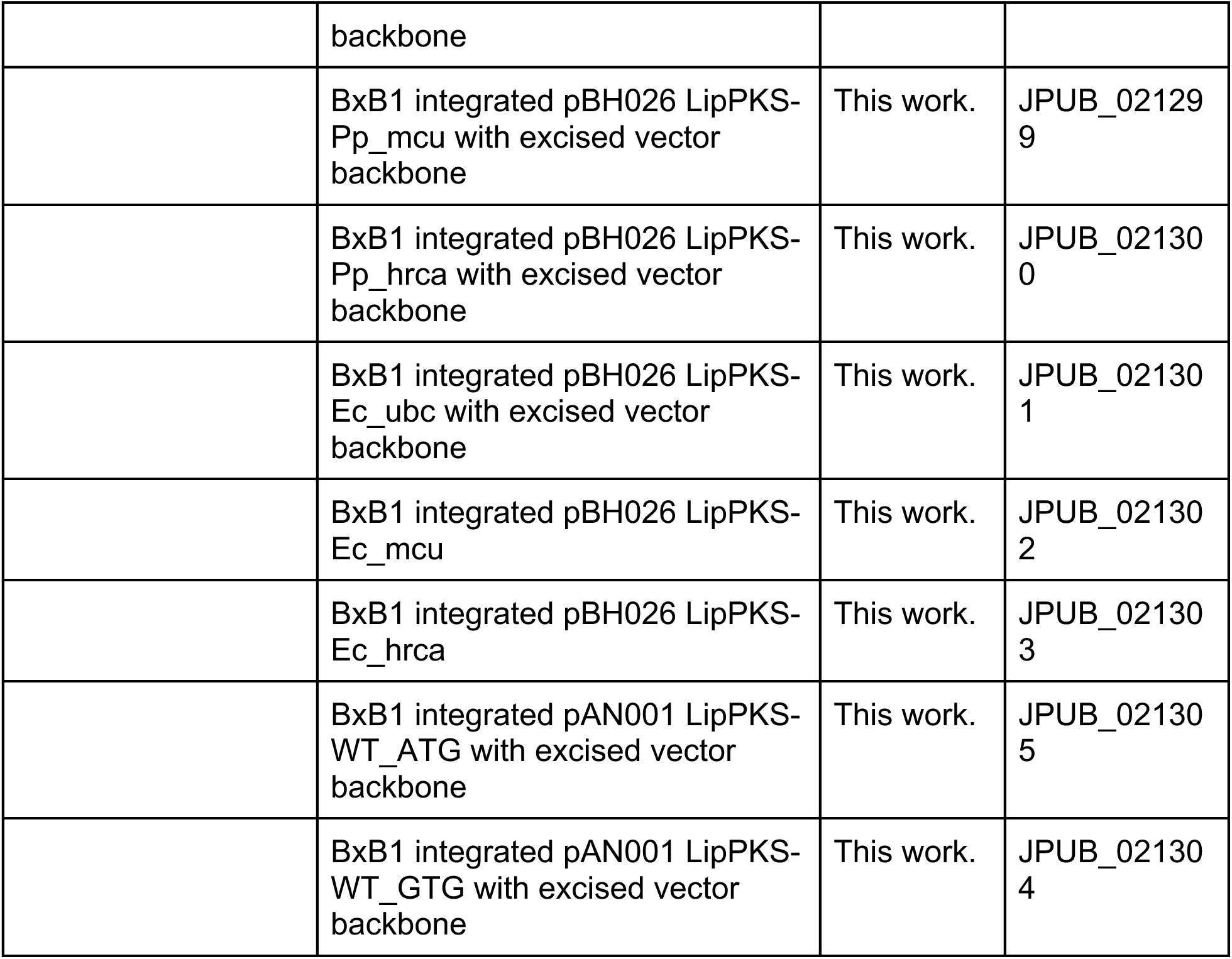
Plasmids and strains used in this study.

Heterologous genes were integrated into the host genome or expressed from plasmid DNA. Integrations of pBH026 or pAN001 vectors and backbone excisions were performed using the SAGE system (22). Excision of the backbones of pBH026 or pAN001 using pGW30 or pALC412 leads to the removal of the repressor *lacI*, the selection marker and origin of replication. Serine recombinase-assisted integrations into the hosts AG5577 and AG6212 were confirmed by colony PCR (cPCR) using primers that flank the specific *attB* site in the host genome and primers that anneal to the 5’ and 3’ end of the target gene.

Engineering of *C. glutamicum* metabolism was performed by first inoculating EPO medium (37 g BHI powder, 25 g glycine, 10 mL Tween, 4 g isoniazid) with a BHI overnight culture. Cells were then grown for 5-6 hours until reaching an optical density (OD) of approximately 1 (λ = 600 nm). Following growth, the cells were washed three times with ice-cold 10% glycerol and then resuspended in 1 mL of ice-cold 10% glycerol. From this suspension, 80 µl of competent cells were transferred to ice-cold electroporation cuvettes with a 2 mm gap. Subsequently, 500 ng of plasmid DNA was used for transformation.

After electroporation, cells were resuspended in 1 mL of BHI medium and heat shocked at 46 ℃ for 5 minutes. Next, cells were recovered at 30℃ for 1-2 hours and cultures were plated on BHI agar supplemented with 25 µg/mL kanamycin and incubated for 2 days. Single colonies were streaked on BHI agar containing 10% sucrose and incubated at 30 ℃ for 1-2 days to allow for the removal of the selection marker. Finally, individual colonies were picked and verified using colony PCR (Table S2).

### 4.4 Plate reader assay for RFP measurement

Cell cultures were harvested after growing for 24 h at 30 and 37 °C, respectively. After centrifuging 500 µL of sample, the cell pellets were washed with 500 µL 0.9 % NaCl solution. OD and fluorescence measurements were performed in a Biotek Synergy H1M plate reader (BioTek, USA) using a black 96-well plate (Corning, USA) and 100 µL sample volume. The OD was measured at 600 nm. The fluorescence of RFP was excited at 535 nm and emission was measured at 620 nm.

### 4.5 Codon optimization of engineered PKS

Codon optimizations were performed using DNA Chisel and the Kazusa database. The target hosts were *E. coli* str. K-12 substr. W3110 (NCBI:txid316407), *C. glutamicum* ATCC13032 (NCBI:txid196627) and *P. putida* KT2440 (NCBI:txid160488). For the hrca method, the LipPKS part was harmonized with *S. aureofaciens* (NCBI:txid1894) and the EryPKS part with *S. erythraea* (NCBI:txid1836). To ensure reproducible results, the Numpy random generator was set to 123. A list of all the applied constraints can be found in the Supplementary (Table S1).

The website for the GUI based on DNA Chisel (https://basebuddy.lbl.gov) was built using the open-source app framework Streamlit. To view the associated files and scripts, or to run a local version of BaseBuddy, see the Github repository (https://github.com/jbei/basebuddy).

### 4.6 Proteomics analysis

Strains for sample preparation were grown at 30 and 37 °C, respectively. After 48 h, 1 mL of cell culture was collected and consolidated in a 96-well plate. Next, the supernatant was removed by centrifuging the plates at 4000 g for 10 min. Cell pellets were then stored at −80 °C until further processing.

Protein was extracted and tryptic peptides were prepared by following established proteomic sample preparation protocol (49). Briefly, cell pellets were resuspended in Qiagen P2 Lysis Buffer (Qiagen, Germany) to promote cell lysis. Proteins were precipitated with addition of 1 mM NaCl and 4 x vol acetone, followed by two additional washes with 80% acetone in water. The recovered protein pellet was homogenized by pipetting mixing with 100 mM ammonium bicarbonate in 20% methanol. Protein concentration was determined by the DC protein assay (BioRad, USA). Protein reduction was accomplished using 5 mM tris 2-(carboxyethyl)phosphine (TCEP) for 30 min at room temperature, and alkylation was performed with 10 mM iodoacetamide (IAM; final concentration) for 30 min at room temperature in the dark. Overnight digestion with trypsin was accomplished with a 1:50 trypsin:total protein ratio. The resulting peptide samples were analyzed on an Agilent 1290 UHPLC system coupled to a Thermo Scientific Orbitrap Exploris 480 mass spectrometer for discovery proteomics (50). Briefly, peptide samples were loaded onto an Ascentis® ES-C18 Column (Sigma–Aldrich, USA) and were eluted from the column by using a 10 minute gradient from 98% solvent A (0.1 % FA in H_2_O) and 2% solvent B (0.1% FA in ACN) to 65% solvent A and 35% solvent B.. Eluting peptides were introduced to the mass spectrometer operating in positive-ion mode and were measured in data-independent acquisition (DIA) mode with a duty cycle of 3 survey scans from m/z 380 to m/z 985 and 45 MS2 scans with precursor isolation width of 13.5 m/z to cover the mass range. DIA raw data files were analyzed by an integrated software suite DIA-NN (51). The databases used in the DIA-NN search (library-free mode) are respective microorganisms’ latest Uniprot proteome FASTA sequences plus the protein sequences of the heterologous proteins and common proteomic contaminants. DIA-NN determines mass tolerances automatically based on first pass analysis of the samples with automated determination of optimal mass accuracies. The retention time extraction window was determined individually for all MS runs analyzed via the automated optimization procedure implemented in DIA-NN. Protein inference was enabled, and the quantification strategy was set to Robust LC = High Accuracy. Output main DIA-NN reports were filtered with a global FDR = 0.01 on both the precursor level and protein group level. The Top3 method, which is the average MS signal response of the three most intense tryptic peptides of each identified protein, was used to plot the quantity of the targeted proteins in the samples (26, 27).

The generated mass spectrometry proteomics data have been deposited to the ProteomeXchange Consortium via the PRIDE partner repository with the dataset identifier PXD042749 (52). DIA-NN is freely available for download from https://github.com/vdemichev/DiaNN.

### 4.7 Polyketide production and quantification by liquid chromatography and mass spectrometry

The conditions used for polyketide production were dependent on the production host. *C. glutamicum* cultures were grown for 96 h, while *P. putida* cells were cultivated for 48 h. Furthermore, the BHI medium used for *C. glutamicum* was supplemented with 5 mM isobutyric acid and 5 mM propionate. The LB medium for *P. putida* contained an additional 20 mM of *L*-valine.

Polyketide production in *E. coli* K207-3 followed the same procedure as described in Yuzawa et al. (2017). In short, cells were grown at 18 °C for 120 h in LB containing 5 mM propionate, kanamycin and 200 µM IPTG.

After cultivation of each host was completed, 300 µL of cell culture was harvested and quenched using 300 µL of −80 °C cold methanol. Next, samples were centrifuged at 14,000 g for 1 min and 300 µL supernatant was transferred into Omega 3K MWCO AcroPrep 96-well filter plates (Pall, USA). After centrifuging for 45 min at 3,000 g, the flow-through was collected and used for further analysis.

LC separation of 3-hydroxy acids was conducted using a Kinetex XB-C18 column (100 mm length, 3 mm internal diameter, 2.6 μm particle size; Phenomenex, USA) with an Agilent 1260 Infinity II LC System (Agilent Technologies, USA) at room temperature. The mobile phase consisted of 0.1% formic acid in water (solvent A) and 0.1% formic acid in methanol (solvent B). The separation of products was achieved with a flow rate of 0.42 mL/min, using the following gradient: 20% to 72.1% B over 6.5 min, 72.1% to 95% B over 1.3 min, and held for 1 min. Subsequently, the flow rate was increased to 0.65 mL/min, and the gradient was 95% to 20% B over 0.2 min, held for 1.2 min.

To identify and quantify the 3-hydroxy acids, the LC system was coupled to an Agilent InfinityLab LC/MSD iQ single quadrupole mass spectrometer (Agilent Technologies, USA), with electrospray ionization (ESI) conducted in the negative-ion mode. Identification of 3-hydroxy acids was performed by comparing the mass and retention time with authentic standards.

### 4.8 Quantitative reverse transcription PCR

For RT-qPCR analysis, samples were collected after 24 h by centrifuging 1 mL cell culture and discarding the supernatant. Total RNA was then extracted using the RNeasy Plus Universal Mini Kit (Qiagen, Germany) following the manufacturer’s instructions. Prior to RNA extraction of *C. glutamicum* samples, cell pellets were treated with 1 mL of 2 mg/mL lysozyme (Roche, Switzerland) and incubated at 30 °C for 10 min. The RNA extracted from the samples was used as a template for complementary DNA (cDNA) synthesis, which was carried out using the LunaScript RT SuperMix Kit (NEB, USA). Subsequently, qPCR was performed using the Luna Universal qPCR Master Mix (NEB, USA). The PCR mixtures were cycled at 95°C for 1 min (one cycle) followed by 40 cycles at 95°C for 15 s and 60°C for 30 s. The amplification profile was monitored on a CFX96 Real-Time PCR Detection System (Bio-Rad, USA). The expression ratio from qPCR was calculated from the Ct value difference between target gene and *rpoD* for *P. putida*, *E. coli* or *rpoC* for *C. glutamicum* as housekeeping genes. Primers used for housekeeping genes and target gene quantification can be found in the Supplementary (Table S2).

### 4.9 Bioinformatic analyses

For phylogenetic analysis, 16S rRNA sequences were extracted from the RNAcentral database (53). The sequences were aligned using the CLUSTAL W algorithm, implemented in MEGA 11 (54). Subsequently, a phylogenetic tree was constructed using the maximum-likelihood method with the Tamura-Nei model.

Codon usage data for organisms part of the Refseq index were obtained from the CoCoPUTS database. For each entry in the database (235025 entries), the codon usage data was normalized by the total number of respective codons analyzed. Taxonomic classifications corresponding to each Refseq entry were retrieved from the National Center for Biotechnology Information (NCBI) database, providing a hierarchical framework for taxonomic inference. To discern the general similarity of codon usage across organisms, a Principal Component Analysis (PCA) was implemented on the normalized codon tables. The first two principal components, accounting for the highest explained variance, were selected and visualized via a scatter plot.

## Acknowledgements

We would like to express our profound appreciation to Adam Guss and his colleagues for generously providing us with the SAGE vector suite, as well as the strains *P. putida* AG5577 and *C. glutamicum* AG6212. Their contributions have been instrumental in advancing our research. Additionally, we want to thank Peter Mellinger and Chenyi Li for their assistance during various stages of the cloning process. This research was funded by the DOE Joint BioEnergy Institute (https://www.jbei.org) supported by the U.S. Department of Energy, Office of Science, Office of Biological and Environmental Research through contract [DE-AC02-05CH11231] between Lawrence Berkeley National Laboratory and the U.S. Department of Energy, by award EE0008926 from the BioEnergy Technologies Office, Office of Energy Efficiency and Renewable Energy, U.S. Department of Energy, and the Philomathia Foundation. A.A.N. was supported by a National Science Foundation Graduate Research Fellowship, fellow ID [2018253421].

## Contributions

Conceptualization: M.S.; Methodology, M.S.; Investigation: M.S., C.Z., N.L., A.A.N., J.B.R., L.K., A.V., C.J.P., Y.C.; Writing – Original Draft: M.S.; Writing – Review and Editing: All authors; Resources and supervision: R.W.H., L.M.B., J.D.K.

