## Supplementary_File_1 for "Maximizing Heterologous Expression of Engineered Type I Polyketide Synthases: Investigating Codon Optimization Strategies"

**Table S1:** List of all applied constraints during codon optimization

| **Codon optimization constraints** |
|---|
| UniquifyAllKmers(9, include_reverse_complement=True) |
| AvoidHairpins(stem_size=10,hairpin_window=100) |
| AvoidPattern("9xA") |
| AvoidPattern("9xT") |
| AvoidPattern("6xC") |
| AvoidPattern("6xG") |
| AvoidPattern("NdeI_site") |
| AvoidPattern("XhoI_site") |
| AvoidPattern("SpeI_site") |
| AvoidPattern("BamHI_site") |
| AvoidPattern("BsaI_site") |
| EnforceGCContent(mini=0.3, maxi=0.75, window=50) |
| EnforceTranslation() |

**Table S2:** Primers used in this study.

| Primer | Sequence | Description |
|---|---|---|
| **Confirmation primers for serine recombinase-assisted genome engineering** | | |
| cPCR-AG5577_BxB1_F | AGCAAATTCGGCAACACGC | Confirmation of BxB1 integrations into AG5577 BxB1 attB site |
| cPCR-AG5577_BxB1_R | CTAGGCAGAATTTTGGGAGTGGC | Confirmation of BxB1 integrations into AG5577 BxB1 attB site |

| | | |
|---|---|---|
| cPCR-AG5577_MR11_F | GTGATTTGAAAGAGTTGTCAGTTAGCTCG | Confirmation of MR11 integrations into AG5577 MR11 attB site |
| cPCR-AG5577_MR11_R | AGGACTCACCTCTAGAACACGC | Confirmation of MR11 integrations into AG5577 MR11 attB site |
| cPCR-AG6212_BxB1_F | CGAATTCTTTCATTTAAGACCCTAATA | Confirmation of BxB1 integrations into AG6212 BxB1 attB site |
| cPCR-AG6212_BxB1_R | CTAGGCAGAATTTTGGGAGTGG | Confirmation of BxB1 integrations into AG6212 BxB1 attB site |
| cPCR-LipPKS-Cg_ubc_F | CGCCATGGAACTGCGTAA | Anneals to 3’ end of codon optimized LipPKS gene. Confirmation of integration into the host genome |
| cPCR-LipPKS-Cg_ubc_R | GGTATGGTGCAAAGGCATCT | Anneals to 5’ end of codon optimized LipPKS gene. Confirmation of integration into the host genome |
| cPCR-LipPKS-Cg_mcu_F | CGCCATGGAATTGCGAAAC | Anneals to 3’ end of codon optimized LipPKS gene. Confirmation of integration into the host genome |
| cPCR-LipPKS-Cg_mcu_R | AAAGACTGTGGCCCAAGTAG | Anneals to 5’ end of codon optimized LipPKS gene. Confirmation of integration into the host genome |
| cPCR-LipPKS-Cg_hrca_F | AGAGCTGGACTCTGGAACT | Anneals to 3’ end of codon optimized LipPKS gene. Confirmation of integration into the host genome |

| | | |
|---|---|---|
| cPCR-LipPKS-Cg_hrca_R | GCGTTCCAAAGGTCGGTTA | Anneals to 5' end of codon optimized LipPKS gene. Confirmation of integration into the host genome |
| cPCR-LipPKS-Pp_ubc_F | CGCCATGGAACTGCGTAA | Anneals to 3' end of codon optimized LipPKS gene. Confirmation of integration into the host genome |
| cPCR-LipPKS-Pp_ubc_R | GTCACGCTATCCAGCAGAC | Anneals to 5' end of codon optimized LipPKS gene. Confirmation of integration into the host genome |
| cPCR-LipPKS-Pp_mcu_F | GCCATGGAACTGCGAAAC | Anneals to 3' end of codon optimized LipPKS gene. Confirmation of integration into the host genome |
| cPCR-LipPKS-Pp_mcu_R | GTCACGGAATCCAGCAGAC | Anneals to 5' end of codon optimized LipPKS gene. Confirmation of integration into the host genome |
| cPCR-LipPKS-Pp_hrca_F | CTTGACCGCCATGGAACT | Anneals to 3' end of codon optimized LipPKS gene. Confirmation of integration into the host genome |
| cPCR-LipPKS-Pp_hrca_R | TCACGCTATCCAGCAGAGA | Anneals to 5' end of codon optimized LipPKS gene. Confirmation of integration into the host genome |
| cPCR-LipPKS-Ec_ubc_F | GCGACGTTGGCTTTGATAG | Anneals to 3' end of codon optimized LipPKS gene. Confirmation of integration into the host genome |

| | | |
|---|---|---|
| cPCR-LipPKS-Ec_ubc_R | CAAATAACGCTGGCTGCAC | Anneals to 5’ end of codon optimized LipPKS gene. Confirmation of integration into the host genome |
| cPCR-LipPKS-Ec_mcu_F | CGCCATGGAACTGCGTAA | Anneals to 3’ end of codon optimized LipPKS gene. Confirmation of integration into the host genome |
| cPCR-LipPKS-Ec_mcu_R | CACGGAATCCAGCAGACT | Anneals to 5’ end of codon optimized LipPKS gene. Confirmation of integration into the host genome |
| cPCR-LipPKS-Ec_hrca_F | TGGGCTTCGACAGCTTAAC | Anneals to 3’ end of codon optimized LipPKS gene. Confirmation of integration into the host genome |
| cPCR-LipPKS-Ec_hrca_R | TCACGCTATCCAGCAGACT | Anneals to 5’ end of codon optimized LipPKS gene. Confirmation of integration into the host genome |
| cPCR-LipPKS-WT_R | CGCAGTCGTGCAGCTTA | Anneals to 5’ end of codon optimized LipPKS gene. Confirmation of integration into the host genome |
| cPCR-LipPKS-WT_F | TCTTCGACCACCCGACA | Anneals to 3’ end of codon optimized LipPKS gene. Confirmation of integration into the host genome |
| cPCR-Sc_mmCoA_F | GTCGGCAAGAGCACGTT | Anneals to 3’ end of S. cellulosum So ce56 mmCoA operon. Confirmation of |

| | | |
|---|---|---|
| | | integration into the host genome |
| cPCR-Sc_mmCoA_R | ATCGTGAAGCGCAGCTC | Anneals to 5’ end of S. cellulosum So ce56 mmCoA operon. Confirmation of integration into the host genome |
| **Confirmation primers for gene deletions and replacements in AG6212** | | |
| cPCR-prpDBC2_F | GTGTTACCGATCCACTGG GCGTCAACG | Confirmation of *prpDBC2* deletion |
| cPCR-prpDBC2_F | CATTGCGCATTCCGATCA TGCGCGTCTGCG | Confirmation of *prpDBC2* deletion |
| cPCR-kivd-CCL4_F | GACATCATTCACCAGCAG GTCGGTGGACTTCGTGC | Confirmation of *kivd-CCL4* replacing Cgl0605 |
| cPCR-kivd-CCL4_R | GCGGTGTGCGGGATGACT TCGGAGTAGATG | Confirmation of *kivd-CCL4* replacing Cgl0605 |
| cPCR-sfp_F | GTCAATGTTGACGCGGCC TGGACGGGTGGAACCGG T | Confirmation of *sfp* replacing Cgl1016 |
| cPCR-sfp_F | ACCATCCGCCACATCGAG TCTGTCCACCAGCT | Confirmation of *sfp* replacing Cgl1016 |
| **Primers for RT-qPCR analysis** | | |
| qPCR-LipPKS-Cg_hrca_F | ATGTCCGAACACAGAGGA AGT | Amplification of LipPKS cDNA for qPCR |
| qPCR-LipPKS-Cg_mcu_F | CATATGTCCGAGCACAGG G | Amplification of LipPKS cDNA for qPCR |
| qPCR-LipPKS-Cg_ubc_F | ATGTCCGAACACAGGGGT AG | Amplification of LipPKS cDNA for qPCR |
| qPCR-LipPKS-Ec_hrca_F | ATGAGCGAACACAGAGGA TCA | Amplification of LipPKS cDNA for qPCR |
| qPCR-LipPKS-Ec_mcu_F | ATGAGCGAACATCGTGGT AGT | Amplification of LipPKS cDNA for qPCR |

| | | |
|---|---|---|
| qPCR-LipPKS-Ec_ubc_F | ATGAGCGAACATCGTGGC | Amplification of LipPKS cDNA for qPCR |
| qPCR-LipPKS-Pp_hrca_F | CATATGAGCGAGCACAGGG | Amplification of LipPKS cDNA for qPCR |
| qPCR-LipPKS-Pp_mcu_F | CATATGTCGGAGCACAGGG | Amplification of LipPKS cDNA for qPCR |
| qPCR-LipPKS-Pp_ubc_F | CATATGAGCGAGCACCGG | Amplification of LipPKS cDNA for qPCR |
| qPCR-LipPKS-WT_GTG_F | CATGTGTCCGAACACCGTG | Amplification of LipPKS cDNA for qPCR |
| qPCR-LipPKS-WT_ATG_F | CATATGTCCGAACACCGTGG | Amplification of LipPKS cDNA for qPCR |
| qPCR-LipPKS-Cg_hrca_R | TCTAAGATGAGTCCTCAGAGCCT | Amplification of LipPKS cDNA for qPCR |
| qPCR-LipPKS-Cg_mcu_R | CTAAGATGCGTTCGCAGAGC | Amplification of LipPKS cDNA for qPCR |
| qPCR-LipPKS-Cg_ubc_R | GAGATGGGTTCGCAGAGCTT | Amplification of LipPKS cDNA for qPCR |
| qPCR-LipPKS-Ec_hrca_R | AGATGAGTTCTCAGAGCCTCG | Amplification of LipPKS cDNA for qPCR |
| qPCR-LipPKS-Ec_mcu_R | GTTCGCAGAGCCTCGCTA | Amplification of LipPKS cDNA for qPCR |
| qPCR-LipPKS-Ec_ubc_R | GTAAATGGGTTCGCAGAGCTTC | Amplification of LipPKS cDNA for qPCR |
| qPCR-LipPKS-Pp_hrca_R | GTTCGCAGAGCCTCGGAT | Amplification of LipPKS cDNA for qPCR |
| qPCR-LipPKS-Pp_mcu_R | GTTCGCAGAGCCTCGCTA | Amplification of LipPKS cDNA for qPCR |
| qPCR-LipPKS-Pp_ubc_R | GTTCGCAGAGCCTCGCTA | Amplification of LipPKS cDNA for qPCR |
| qPCR-LipPKS-WT_GTG_R | AGATGCGTGCGCAGAG | Amplification of LipPKS cDNA for qPCR |

| | | |
|---|---|---|
| qPCR-LipPKS-WT_ATG_R | AGATGCGTGCGCAGAG | Amplification of LipPKS cDNA for qPCR |
| qPCR-Pp-RpoD_F | CAGGGCTACCTGACTTAC GC | *P. putida* housekeeping gene used as internal control for qPCR |
| qPCR-Pp-RpoD_R | ACGTTGATCCCCATGTCG TT | *P. putida* housekeeping gene used as internal control for qPCR |
| qPCR-Ec-RpoD_F | GGAGCAAAACCCGCAGTC | *E. coli* housekeeping gene used as internal control for qPCR |
| qPCR-Ec-RpoD_R | CGACGATATCTTCCGGCA G | *E. coli* housekeeping gene used as internal control for qPCR |
| qPCR-Cg-RpoC_F | GTGCTCGACGTAAACGTC TTC | *C. glutamicum* housekeeping gene used as internal control for qPCR |
| qPCR-Cg-RpoC_R | GAGGGTTCGGTAGTTGAT GGT | *C. glutamicum* housekeeping gene used as internal control for qPCR |
| **Gibson primers for assembly of pAN001 LipPKS-WT_GTG** | | |
| gPCR-WT-LipM1-Frag1_F | CGAATTCAAAAGATCTTT TAAGAAGGAGATATACAT GTGTCCGAACACCGTGG CAGTGC | Amplification of WT sequence of LipM1 fragment 1 |
| gPCR-WT-LipM1-Frag1_R | CCTCGTCGCGAGGGCG AACGCCACGTCGGCG | Amplification of WT sequence of LipM1 fragment 1 |
| gPCR-WT-LipM1-Frag2_F | ACGTGGCGTTCGCCCTC GCGACGAGGCGTACTGC | Amplification of WT sequence of LipM1 fragment 2 |
| gPCR-WT-LipM1-Frag2_R | TCCCGCTGTCCAGCTCG GCCTTGAGGTACGCG | Amplification of WT sequence of LipM1 fragment 2 |

| | | |
|---|---|---|
| gPCR-WT-EryM6-TE_F | CCTCAAGGCCGAGCTGGACAGCGGGACTCCCGCCC | Amplification of WT sequence of EryM6-TE |
| gPCR-WT-EryM6-TE_R | GTTTTATTTGATGCCTGGAGATCCTTACTCGATCAGTGGTGGTGGTGGTGGTGC | Amplification of WT sequence of EryM6-TE |
| **Gibson primers for assembly of pAN001 LipPKS-WT_ATG** | | |
| gPCR-WT-LipM1-ATG-Frag1_F | CGAATTCAAAAGATCTTTTAAGAAGGAGATATACATATGTCCGAACACCGTGGCAGTGC | Amplification of WT sequence of LipM1 fragment 1 with ATG start codon mutation |
| **Gibson primers for assembly of pBH026 RFP** | | |
| gPCR-pBH026_F | GAAGGTCGTCACTCCACCGGTGCTTAAGGATCCAAACTCGAGTAAGGATCTCCAGG | Amplification of pBH026 plasmid backbone |
| gPCR-pBH026_R | ACGTCTTCGCTACTCGCCATATGTATATCTCCTTCTTAAAAGATCTTTTGAATTCG | Amplification of pBH026 plasmid backbone |
| gPCR-RFP_F | AGAAGGAGATATACATATGGCGAGTAGCGAAGACG | Amplification of *rfp* gene |
| gPCR-RFP_R | TTAAGCACCGGTGGAGTGACGACCT | Amplification of *rfp* gene |
| **Gibson primers for assembly of pK18 ΔCgl0605::kivd-CCL4** | | |
| gPCR-pK18-Cgl0605_F | TCGGGTGGGCCTTTCTGCGTTTATAGCCCTTGATTATTGCCAAAGAAACCTTTAAGGACT | Amplification of pK18 ΔCgl0605 plasmid backbone |
| gPCR-pK18-Cgl0605_R | CGAGCCGCAGCCGAATGTGACTAGTTAACGTGCAGGCTTACCTTTTGGAAGC | Amplification of pK18 ΔCgl0605 plasmid backbone |

| | | |
|---|---|---|
| gPCR-kivd-CCL4_F | GCTTCCAAAAGGTAAGCC TGCACGTTAACTAGTCAC ATTCGGCTGCGGCTCG | Amplification of *kivd* and *CCL4* expression cassette |
| gPCR-kivd-CCL4_R | AGTCCTTAAAGGTTTCTTT GGCAATAATCAAGGGCTA TAAACGCAGAAAGGCCCA CCCGA | Amplification of *kivd* and *CCL4* expression cassette |
| **Gibson primers for assembly of pJH209 Sc_mmCoA** | | |
| gPCR-pJH209_F | GGCCAGGAACCGTAAAAA AGTCAAAAGCCTCCGGTC GGAGG | Amplification of pJH209 plasmid backbone |
| gPCR-pJH209_R | CGACTTCGTGACAACGAT CCCCCAACTGAGAGAACT CAAAGG | Amplification of pJH209 plasmid backbone |
| gPCR-Ptac-Sc_mmCoA_F | AGTTCTCTCAGTTGGGGG ATCGTTGTCACGAAGTCG ACTACG | Amplification of *S. cellulosum* So56 mmCoA operon under the control of $P_{tac}$ |
| gPCR-Ptac-Sc_mmCoA_F | GACCGGAGGCTTTTGACT TTTTTACGGTTCCTGGCCT TTTGC | Amplification of *S. cellulosum* So56 mmCoA operon under the control of $P_{tac}$ |
| **Gibson primers for assembly of pGingerBG-NahR LipPKS-Pp_mcu** | | |
| gPCR-pGingerBG-NahR_F | GCGGCAATAGCTGAGGA TCCAAACTCGAGTAAGG ATCTCC | Amplification of pGingerBG-NahR plasmid backbone |
| gPCR-pGingerBG-NahR_R | ACCCCTGTGCTCCGACA TATGTATATCTCCTTCTT AAATGATGGCTTTATTG | Amplification of pGingerBG-NahR plasmid backbone |
| gPCR-LipPKS-Pp_mcu-Frag1_F | GAAGGAGATATACATAT GTCGGAGCACAGGGGTA GTG | Amplification of LipPKS-Pp_mcu fragment 1 |
| gPCR-LipPKS-Pp_mcu-Frag1_R | CGCAAGCCTGCCATAGT GCGGCCAGACTCACC | Amplification of LipPKS-Pp_mcu fragment 1 |

| | | |
|---|---|---|
| gPCR-LipPKS-Pp_mcu-Frag2_F | TCTGGCCGCACTATGGCAGGCTTGCGGGGTG | Amplification of LipPKS-Pp_mcu fragment 2 |
| gPCR-LipPKS-Pp_mcu-Frag2_R | CGAGTTTGGATCCTCAGCTATTGCCGCCGCCCAG | Amplification of LipPKS-Pp_mcu fragment 2 |
| **Gibson primers for assembly of pGingerBG-NahR LipPKS-Ec_mcu** | | |
| gPCR-pGingerBG-NahR_F | GGCGGAGGGAATAGTTAAGGATCCAAACTCGAGTAAGGATCTCC | Amplification of pGingerBG-NahR plasmid backbone |
| gPCR-pGingerBG-NahR_R | CCACGATGTTCGCTCATATGTATATCTCCTTCTTAAATGATGGCTTTATTG | Amplification of pGingerBG-NahR plasmid backbone |
| gPCR-LipPKS-Ec_mcu-Frag1_F | GAAGGAGATATACATATGAGCGAACATCGTGGTAGTGCAGGT | Amplification of LipPKS-Ec_mcu fragment 1 |
| gPCR-LipPKS-Ec_mcu-Frag1_R | GTTCTTCATCACCGCTACGTAGAACCTCCAGCAGGTTCCAATCCA | Amplification of LipPKS-Ec_mcu fragment 1 |
| gPCR-LipPKS-Ec_mcu-Frag2_F | GGAGGTTCTACGTAGCGGTGATGAAGAACTGAGTAATCGCG | Amplification of LipPKS-Ec_mcu fragment 2 |
| gPCR-LipPKS-Ec_mcu-Frag2_R | CGAGTTTGGATCCTTAACTATTCCCTCCGCCCAGCC | Amplification of LipPKS-Ec_mcu fragment 2 |

36

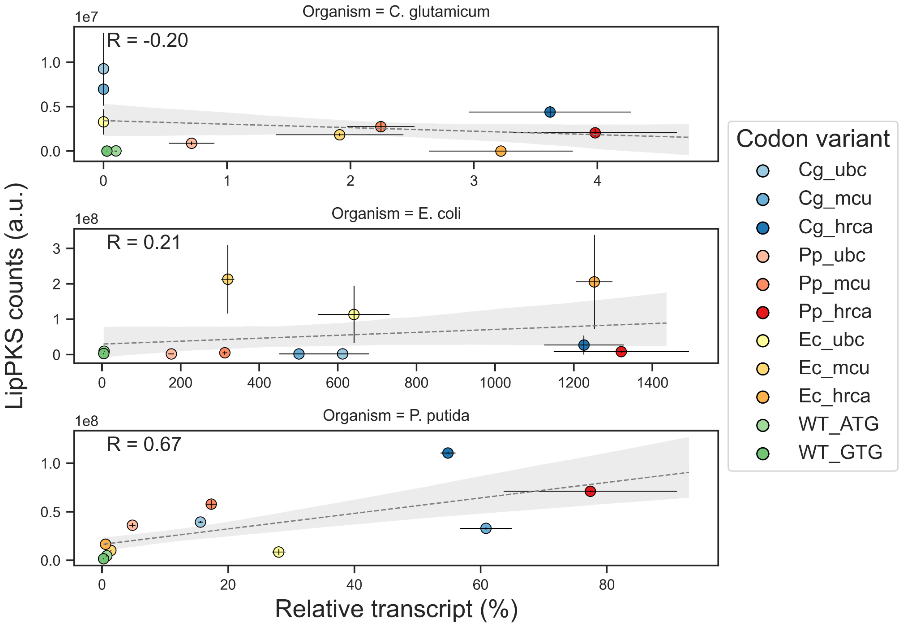


**Figure S1:** Regression plot of LipPKS counts and relative transcript for *C. glutamicum*, *E. coli*, and *P. putida* (n = 3). Only *P. putida* showed a slight correlation between these two parameters.

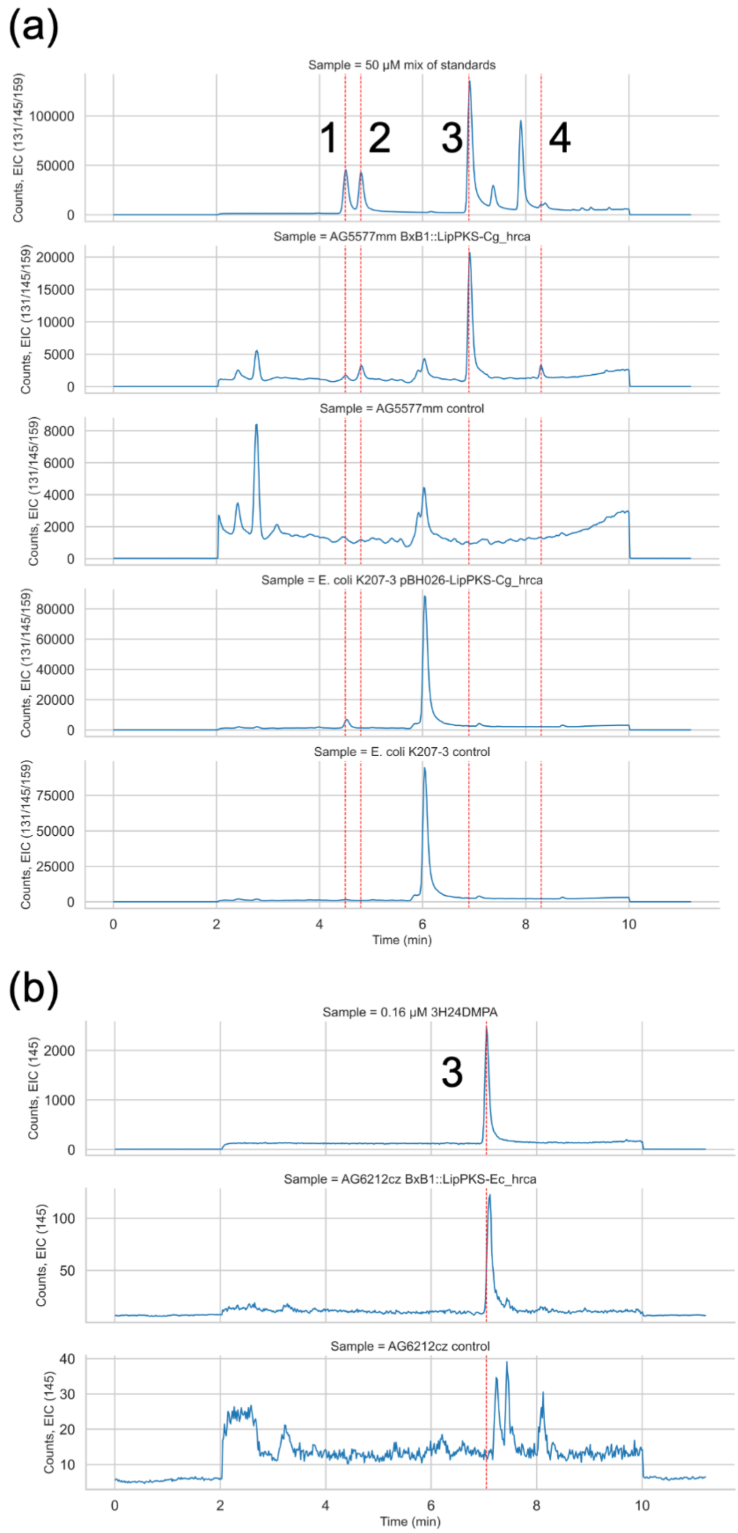


**Figure S2:** LC-MS chromatograms showing the production of unnatural polyketides in *C. glutamicum* (a), *E. coli* and *P. putida* (b). Red lines are indicating the retention time of the authentic standard. Additional peaks for the authentic standards are most likely caused by racemic mixtures of the 3-hydroxy acids. Standards for (3R,2R)-3-hydroxy-2,4-dimethylpentanoic acid (3H24DMPA) and (3R,2R)-3-hydroxy-2-methylpentanoic acid (3H2MPA) are enantiopure. Standards for 3-hydroxy-4-methylpentanoic acid (3H4MPA) and 3-hydroxy-2,4-dimethylhexanoic acid (3H24DMHA) are racemic mixtures. 1 = 3H2MPA; 2 = 3H4MPA; 3 = 3H24DMPA; 4 = 3H24DMHA

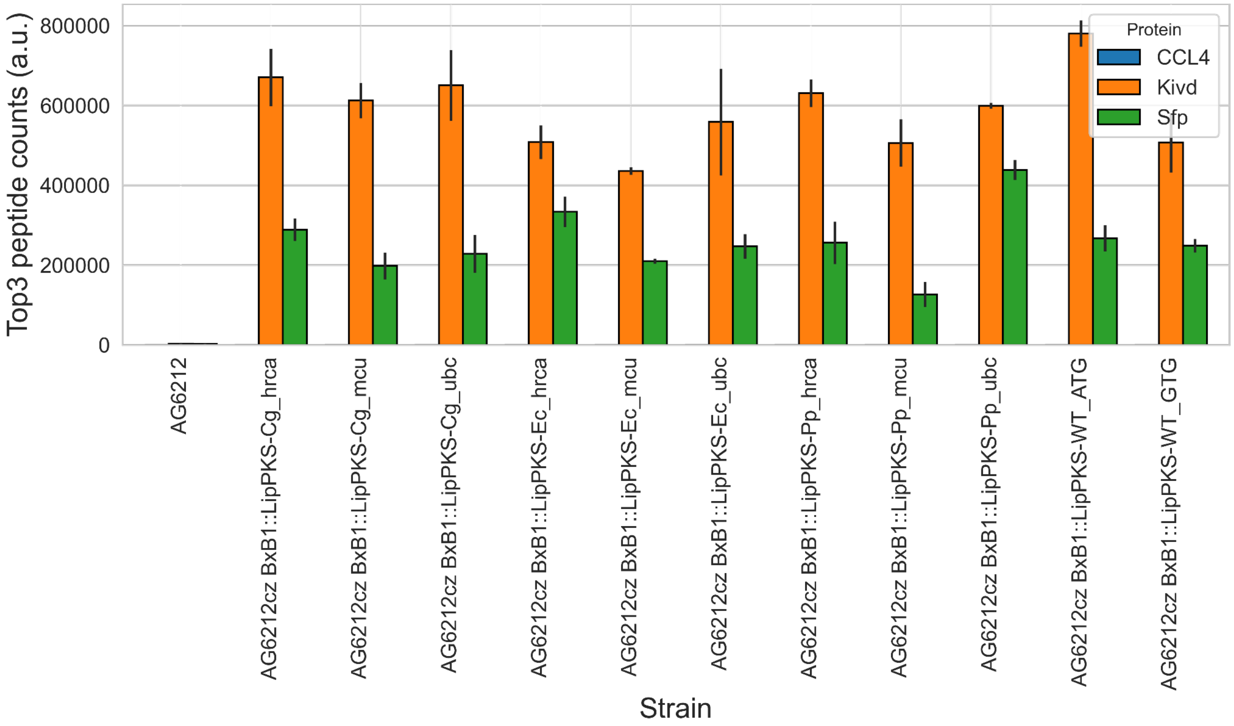


**Figure S3:** Detection of peptides for supplementary pathways in the *C. glutamicum* ATCC 13032 derivative AG6212cz (n = 3). The corresponding peptides for the phosphopantetheinyl transferase (PPTase) Sfp and ketoisovalerate decarboxylase, Kivd, were detected in all AG6212cz strains. CCL4 peptides could not be detected.

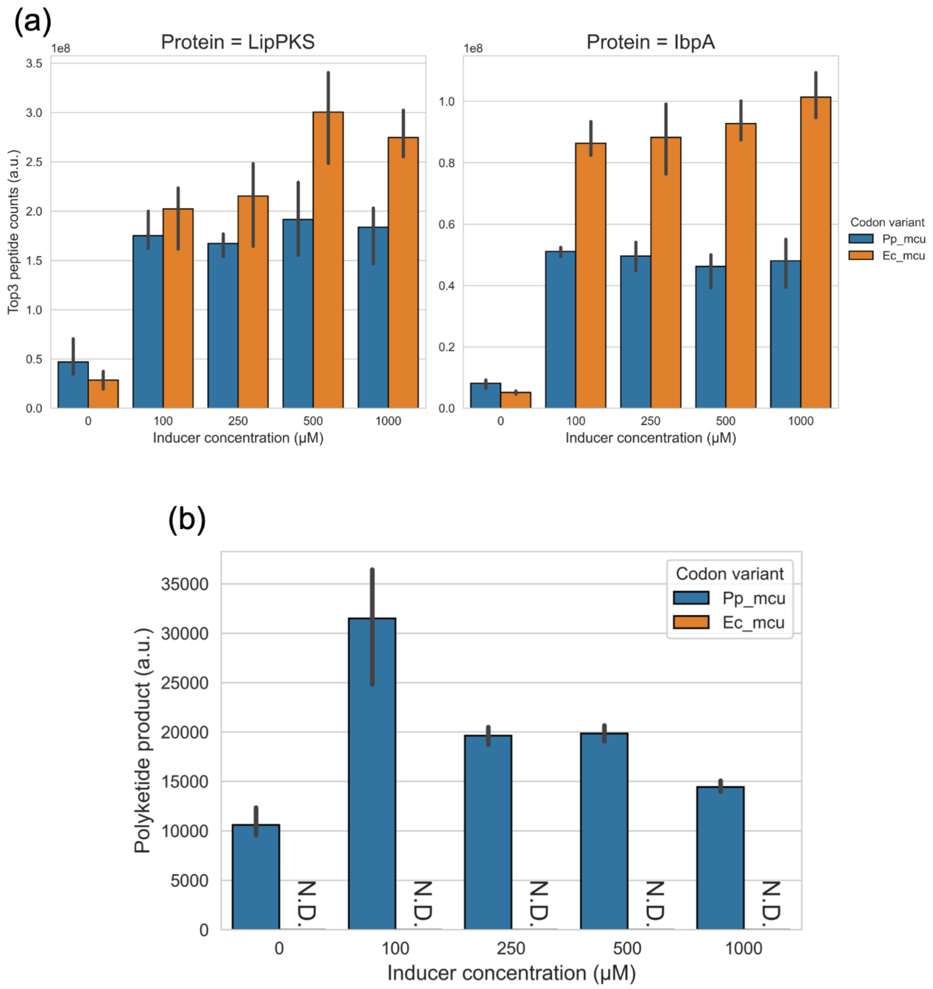


**Figure S4:** Expression of the LipPKS codon variants Pp_mcu and Ec_mcu from the inducible vector system pGingerBG-NahR in *P. putida* (n = 3). The inducer salicylate was added at the time of inoculation. (a) LipPKS protein levels were comparable, while IbpA levels were significantly higher for Ec_mcu. (b) LC-MS analysis of the same samples (n = 3) revealed no production of the corresponding polyketide for the Ec_mcu variant.
